# A proteomics analysis of 5xFAD mouse brain regions reveals the lysosome-associated protein Arl8b as a candidate biomarker for Alzheimer’s disease

**DOI:** 10.1101/2023.01.16.523715

**Authors:** Annett Boeddrich, Christian Haenig, Nancy Neuendorf, Eric Blanc, Andranik Ivanov, Marieluise Kirchner, Philipp Schleumann, Irem Bayraktaroğlu, Matthias Richter, Christine Mirjam Molenda, Anje Sporbert, Martina Zenkner, Sigrid Schnoegl, Christin Suenkel, Luisa-Sophie Schneider, Agnieszka Rybak-Wolf, Bianca Kochnowsky, Lauren M. Byrne, Edward J. Wild, Jørgen E. Nielsen, Gunnar Dittmar, Oliver Peters, Dieter Beule, Erich E. Wanker

## Abstract

**Background:** Alzheimer’s disease (AD) is characterized by the accumulation of amyloid-β (Aβ) peptides in intra- and extracellular deposits. How Aβ aggregates perturb the proteome in brains of patients and AD transgenic mouse models, however, remains largely unclear. State-of-the-art mass spectrometry (MS) methods can comprehensively detect proteomic alterations in neurodegenerative disorders, providing relevant insights unobtainable with transcriptomics investigations. Analyses of the relationship between progressive Aβ aggregation and protein abundance changes in brains of 5xFAD transgenic mice have not been reported previously.

**Methods:** We quantified progressive Aβ aggregation in hippocampus and cortex of 5xFAD mice and controls with immunohistochemistry and biochemical membrane filter assays. Protein changes in different mouse tissues were analysed by MS-based proteomics using label-free quantification (LFQ); resulting MS data were processed using an established pipeline. Results were contrasted with existing proteomic data sets from postmortem AD patient brains. Finally, abundance changes in the candidate marker Arl8b were validated in CSF from AD patients and controls using ELISAs.

**Results:** Experiments revealed a more rapid accumulation of Aβ42 peptides in hippocampus than in cortex of 5xFAD mice, accompanied by many more protein abundance changes in hippocampus than in cortex, indicating that Aβ42 aggregate deposition is associated with brain region-specific proteome perturbations. Generating time-resolved data sets, we defined Aβ aggregate-correlated and anticorrelated proteome changes, a fraction of which was conserved in postmortem AD patient brain tissue, suggesting that proteome changes in 5xFAD mice mimic disease relevant changes in human AD. We detected a positive correlation between Aβ42 aggregate deposition in the hippocampus of 5xFAD mice and the abundance of the lysosome-associated small GTPase Arl8b, which accumulated together with axonal lysosomal membranes in close proximity of extracellular Aβ plaques in 5xFAD brains. Abnormal aggregation of Arl8b was observed in AD brain tissue. Arl8b protein levels were significantly increased in cerebrospinal fluid (CSF) of AD patients, a clinically accessible body fluid.

**Conclusions:** We report a comprehensive biochemical and proteomic investigation of hippocampal and cortical brain tissue derived from 5xFAD transgenic mice, providing a valuable resource to the neuroscientific community. We identified Arl8b, with significant abundance changes in 5xFAD and AD patient brains. Arl8b might enable the measurement of progressive lysosome accumulation in AD patients and have clinical utility as a candidate biomarker.

Data are available via ProteomeXchange with identifier PXD030348.

## Background

Alzheimer’s disease (AD) is the most common form of dementia in the elderly; its prevalence is growing rapidly in aging societies [1]. In 2050, ∼150 million people worldwide are estimated to be affected with AD, imposing a substantial burden on patients and health care systems [1].

AD is a chronic neurodegenerative illness with a long preclinical phase (∼20 years) and an average clinical duration of 8-10 years [2, 3]. Most patients with AD (>95%) have the sporadic form, which is characterized by late disease onset (80-90 years) and progressive neurodegeneration in wide areas of the cerebral cortex and the hippocampus [3, 4]. Sporadic AD is also often associated with the accumulation of amyloid-β (Aβ) peptides in intracellular deposits and extracellular plaques [5] as well as the aggregation of the microtubule protein tau, leading to the formation of neurofibrillary tangles (NFTs) in neurons [6]. Aβ peptides are derived by proteolytic cleavage of the amyloid precursor protein (APP) by γ- and β-secretases, which include presenilin 1 (PS1) and presenilin 2 (PS2) that are encoded by the genes *PSEN1* and *PSEN2*, respectively [7, 8].

The “amyloid cascade hypothesis” states that the release of Aβ peptides from APP and their self-assembly into amyloidogenic aggregates are the direct cause of dementia in AD patients [9, 10]. Thus, the formation of “proteopathic” Aβ assemblies are potentially at the root of AD and may drive all subsequent molecular pathogenic processes that eventually lead to appearance of disease symptoms such as memory loss and progressive cognitive impairment. However, the linearity of the Aβ-driven disease cascade remains controversial. It provides no good explanation for the very long prodromal phase in AD as well as for the relatively weak correlation between the abundance of amyloid plaques in specific brain regions and the much later observed disease symptoms in sporadic AD patients [11].

A more direct causal link between Aβ production and disease development is established for familial AD (FAD), where causative mutations in *APP*, *PSEN1* and *PSEN2* lead to early onset of AD (mean age of ∼45 years) and the enhanced production of a 42-residue-long Aβ peptide (Aβ42) that is a major component of extracellular amyloid plaques in patient brains [12] and rapidly aggregates in cell-free assays and AD models [13]. Experimental evidence was obtained that imbalance between production and clearance of Aβ42 and related Aβ peptides is a very early, even initiating event in AD [14]. This is also supported by the discovery of AD risk proteins such as ApoE4, SORL1 and PICALM that are all thought to play a specific functional role in Aβ clearance in AD brains [15–17]. Soluble Aβ42 oligomers isolated directly from cortex of patients can dose-dependently decrease synaptic function and significantly impair memory of a learned behaviour in healthy adult rats, representing further evidence for disease relevance of the polypeptide [18]. Finally, biomarker studies with FAD patients carrying *APP*, *PSEN1* and *PSEN2* mutations have elucidated the pathogenic sequence of events in AD brains. In this context, it was demonstrated that the Aβ42 levels in CSF start to decrease ∼25 years before expected symptom onset [2]. This is then followed by the appearance of fibrillar PIB-reactive amyloid deposits (detected by PET studies), increased levels of tau in CSF and progressive brain atrophy roughly 15 years before expected symptom onset [2]. Thus, it seems adequate to speculate that progressively accumulating Aβ42 peptides cause “aggregate” stress in FAD patient brains [19], which finally leads to progressive neurodegeneration and dementia.

The 5xFAD transgenic (tg) mouse model recapitulates major features of AD amyloid pathology [20, 21]. These mice express both human APP and PS1 with five FAD mutations in neurons, leading to overproduction of Aβ42 peptides, which progressively accumulate in intraneuronal deposits and extracellular amyloid plaques [5] in mouse brains. Strikingly, Aβ42 aggregation in 5xFAD mouse brains is accompanied by activated neuroinflammation, loss of synapse functions and neurodegeneration, which are all well-known pathobiological features of AD patient brains [20, 22–24]. In a recent proteomics study, convincing data were obtained that proteome changes in brains of 5xFAD tg mice and AD patients are cross-correlated [25], supporting the hypothesis that this disease model mimics key aspects of AD pathogenesis.

Here, we have investigated the impact of progressive Aβ42 aggregation on the proteome in hippocampus and cortex of 5xFAD tg mice to elucidate the direct molecular consequences of Aβ42-aggregate stress in AD mouse brains. We hypothesized that the proteome response to Aβ42 aggregates might be distinct in hippocampus and cortex, because protein composition and abundance are significantly different in these brain regions [26]. Previous ‘OMICS’ studies have shown changes in transcript and protein levels in brains of 5xFAD tg mice [20, 22–25, 27, 28]. However, a direct link between progressive Aβ42 accumulation and proteome changes in specific disease-relevant brain regions such as hippocampus and cortex remains to be elucidated. Also, the concordance of proteome changes in in 5xFAD and AD patient brains has previously not been assessed.

Here, we present experimental results indicating that progressive Aβ42-mediated aggregate stress is associated with brain region-specific proteome changes in 5xFAD mouse brains. We observed that Aβ42 aggregation was more pronounced in hippocampus than in cortex in AD mouse brains and that this phenomenon was also accompanied by a high number of proteins that changed in their abundance. Utilizing our data sets, we defined Aβ aggregate-correlated and anticorrelated proteome changes in 5xFAD brains, a fraction of which was also conserved in postmortem AD patient brains, suggesting that the observed proteome changes in mouse brains at least in part reflect the pathogenic process in patient brains. Furthermore, our investigations revealed that the small GTPase Arl8b, which controls lysosomal-related vesicle transport in neurons [29], strongly correlates with the progressive accumulation of Aβ42 aggregates and is abnormally enriched in the vicinity of amyloid plaques in brains of 5xFAD tg mice. Finally, elevated levels of Arl8b were found in postmortem brain and CSF samples of AD patients, suggesting that this protein is of disease relevance and a candidate AD biomarker. The implications of our results for the development of protein biomarkers that potentially monitor the abnormal accumulation of lysosomes in response to Aβ aggregation in AD patient brains are discussed.

## Methods

### Antibodies

Commercially available antibodies applied in this study are shown in **Additional File 1: Table S1**. The concentrations of antibodies used for immunoblotting (IB), ELISA and immunostaining (IS) of brain slices are described in the method sections below. Secondary Alexa Fluor-labelled anti-rat, anti-mouse and anti-rabbit antibodies were used for immunostaining, while peroxidase labelled anti-rabbit and anti-mouse secondary antibodies were used for immunoblotting. Alexa Fluor 594-labelled anti-β-Amyloid 1-16 (clone 6E10) antibody is a fluorophore-labelled primary antibody which was used for immunostaining of amyloid-β (Aβ). A labelling with a secondary antibody is not necessary in this case. Production and characterization of the mouse monoclonal antibody 352 applied for immunostaining was described previously [30]. This antibody specifically recognizes fibrillar Aβ42 aggregates but not monomers in dot blot assays.

### Analysis of CSF samples of AD and HD patients

AD and control CSF samples (**Additional file 1: Table S2**) were obtained from patients at the Memory Clinic of Charité Universitätsmedizin Berlin. The procedures for CSF sample collection and treatment were described elsewhere [31, 32]. Briefly, CSF was collected in polypropylene tubes. Immediately after collection, the tubes were gently shaken and centrifuged (2,000 x g; room temperature; 10 min).

Supernatant was taken off, aliquoted, frozen in liquid nitrogen and stored at -80 °C. To quantify Aβ peptides in CSF, the Lumipulse^®^ G β-Amyloid 1-42 and Lumipulse^®^ G β-Amyloid 1-40 assays (Fujirebio Germany GmbH, Hannover, Germany) were used. For total (t-)tau and phosphorylated (p-)tau quantification, the Lumipulse^®^ G Total Tau and Lumipulse^®^ G pTau 181 assays (Fujirebio Germany GmbH, Hannover, Germany) were used, respectively. Under these conditions, the following CSF biomarker values were rated as indicative of AD: Aβ(1-42) < 680 pg/ml, Aβ(1-42) / Aβ(1-40) ratio <0.055, t-tau >400 pg/ml and p-tau >62 pg/ml (see **Additional file 1: Table S2**). Individuals who did not meet the above-mentioned criteria were defined as control. The control group comprised CSF samples derived from subjective cognitive impairment patients, patients suffering from depression and healthy individuals. The AD group comprised CSF samples from AD patients with different grades of dementia. CSF samples from HD patients and controls were obtained from the Danish Dementia Research Centre [33] and the UCL Huntington’s Disease Centre, University College London, London, UK [34, 35].

### Mouse model and breeding

5xFAD mice B6SJL-Tg(APPSwFlLon, PSEN1*M146L*L286V)6799Vas/Mmjax [21] overexpress two human AD-related proteins: the mutant human APP (695) protein with the Swedish (K670N), Florida (I716V) and the London (V717I) mutations and the human PS1 protein with the mutations M146L and L286V. 5xFAD mice were backcrossed to C57BL/6NCrl mice for more than 10 generations (T. Willnow, MDC Berlin-Buch). For our experiment, female transgenic mice were mated to male wild-type mice. Resulting offspring were weaned at approximately three weeks of age; genotyping for the *APP* and *PSEN1* transgenes was performed by PCR with tissue biopsies [21]. Male mice of 2, 5, and 8 months were used for the experiments. Animal care was in accordance with the directive 2010/63/EU. The MDC has signed the Basel Declaration in 2012 and observes an internal animal welfare directive.

Experimental procedures were approved by the local animal welfare authority in Berlin, Germany under license number TVV G0073/17. Mice were group-housed in cages with wooden bedding, environmental enrichment and *ad libitum* food and water supply.

### Mouse brain tissue processing

Mice, whose brains were used for immunohistochemistry analysis, were transcardially perfused with 25 ml 0.9% saline followed by 90 ml 4% paraformaldehyde in 0.1 M phosphate buffer pH 7.4 (PB) under deep anesthesia using pentobarbital (100-150 mg/kg). Brains were removed rapidly and post-fixed by immersion in 4% PFA in PB overnight. Then, the brains were cryoprotected by incubation in 20% sucrose in PB at 4 °C for 24 h followed by incubation in 30% sucrose in PB at 4 °C for 24h. Brains were divided into hemispheres, frozen in isopentane and stored at -80 °C until cryosectioning. For all other analyses mice were sacrificed by cervical dislocation. Brains were removed and washed in 1xPBS. Hippocampus and cortex were carefully dissected, frozen in liquid nitrogen and stored at -80 °C until further use.

### Preparation of brain homogenates

For preparation of mouse brain homogenates for membrane filter assays (MFAs) and gel analysis, ∼80 mg brain was homogenized in 640 µl ice-cold 50 mM Tris pH 7.5 buffer containing protease inhibitors and Benzonase (0.25 U/µl) using the Precellys homogenizer (CK14 tubes, 5,000 rpm, 14 sec for cortex and CK14 tubes, 4,500 rpm, 2 sec for hippocampus). Then, 160 µl of a concentrated buffer solution containing 250 mM Tris pH 7.5, 750 mM NaCl, 0.5% SDS, 2% sodium deoxycholate and 5% Triton were added to the homogenate and incubated for 30 min at 4 °C on a rotating wheel. The homogenate was transferred to a protein LoBind tube and centrifuged at 1.500 x g for 20 min at 4 °C. Then, the supernatant was carefully removed and pipetted to a new tube. Protein concentration was determined using the Pierce BCA assay (ThermoFisher Scientific, Waltham, MA, USA). Supernatant was stored at - 80 °C until further use. Human postmortem brain tissue samples derived from temporal cortex were obtained from the Newcastle Brain Tissue Resource (NBTR), Newcastle University, UK. The method for preparation of human brain homogenates was described previously [36].

### Quantification of Aβ peptides with ELISA

To determine the amount of human Aβ40 and Aβ42 peptides in mouse brain homogenates, commercial ELISA kits (ThermoFisher Scientific, Waltham, MA, USA: KHB3442, KHB3482) were used according to manufacturer’s instructions. Briefly, mouse brain material was weighed and homogenized using Precellys CK14 tubes as described above. To solubilize Aβ, homogenates were incubated with guanidine hydrochloride (final concentration 5 M) for 3.5 h with 800 rpm shaking at 22 °C. Then, samples were diluted and analysed by ELISA.

### Arl8b ELISA

Human Arl8b protein abundance was determined in CSF samples derived from AD and HD patients as well as control individuals. CSF was centrifuged at 2,000g for 10 min. The supernatant was carefully removed, diluted 1:18 in standard diluent delivered with the Arl8b ELISA kit (CUSABIO, CSB-EL002100HU). All steps were performed according to supplier protocols. Arl8b protein amounts in CSF samples were calculated using an Arl8b ELISA standard curve.

### Western blot analysis

Protein extracts were boiled with NuPAGE 4x sample buffer containing 50 mM DTT for 5 min and then loaded onto NuPAGE Novex 4-12% Bis-Tris gels (ThermoFisher Scientific). Electrophoresis was performed according to a standard protocol followed by transfer of proteins onto PVDF membrane (pore size 0.2 µm, MerckMillipore) using a semidry blotting system (Power Blotter XL, Thermofisher Scientific) and a 2x NuPAGE™ transfer buffer containing 10% methanol. Immunoblotting was performed using the method described previously [30]. The generated blots were blocked for 30 min with 3% skim milk (Sigma-Aldrich) in 1x PBS (13.7 mM NaCl, 0.27 mM KCl, 1 mM Na_2_HPO_4_, 0.2 mM KH_2_PO_4_, pH 7.4) containing 0.05% Tween 20 (PBS-T). Then, the membrane was incubated over night with the primary antibody diluted in 3% skim milk PBS-T (6E10, 1:500; anti-Alpha-Tubulin (#T6074), 1:8,000; anti-Alpha-Tubulin (SAB3501072), 1:2,000; anti-Presenilin-1, 1:500; anti-Arl8b, 1:500; anti-LAMP1, 1:500; anti-Calnexin, 1:1000; anti-VDAC, 1:1000; anti-Golgin97, 1:750; anti-Flotillin, 1:1000, anti-NDUFB3, 1:1000). Subsequently, the membrane was washed three times for 10 min in PBS-T and incubated with the secondary peroxidase-conjugated anti-mouse or anti-rabbit antibody (1:2,000) for 1 h at room temperature. The membrane was washed two times for 5 min in PBS-T and two times for 10 min in PBS; immunoreactive protein was detected using ChemiGlow (Biozym). Unless otherwise mentioned, chemiluminescence was measured with a FujiFilm LAS-3000 and images were quantified using the Aida image analysis software (Raytest).

### Membran filter assay

Mouse cortical and hippocampal brain homogenates were filtered through a cellulose acetate membrane (10 µg per dot) with a pore size of 0.2 µm (GE Healthcare Life Sciences, Munich, Germany). Per cavity of filtration unit, 10 µg homogenate was filtered. Then, the membrane was washed with 1x PBS and blocked for 30 min with 3% skim milk in PBS-T. The protein aggregates retained on filter membranes were finally detected by antibody-based reactions as described for western blotting. The membrane filter assays procedure with human brain homogenates was described previously [36]. Per cavity, 10 µg of homogenate was filtered.

### Cryosectioning

Frozen hemispheres of mouse brains were embedded in Tissue-Tek O.C.T. (Sakura) and sectioned using a CM3050 S Cryostat (Leica). 40 μm thick sections were collected in 24 well plates and stored in cryoprotectant solution (0.65 g NaH_2_PO_4_ × H_2_O, 2.8 g Na_2_HPO_4_ in 250 ml ddH_2_O, pH 7.4 with 150 ml ethylene glycol, 125 ml glycerine) at 4 °C until further processing. Eight sections between bregma -1.4 mm to -3.6 mm [37] at equal distance from each other, determined by Hämalaun/Eosin staining, were used for further analysis.

### Hämalaun/Eosin staining

Free floating sections were washed 2 x 10 min with PBS, mounted on microscope slides and air dried. Hämalaun/Eosin staining was carried out at room temperature. Sections were incubated with Mayer’s Hämalaun (Roth) for 15 min followed by 10 min bluing in tap water. Sections were incubated in Eosin (0.5% solution with a few drops of acetic acid) for 30 sec followed by a wash in H_2_O. Sections were subsequently treated with 50, 70 and 99% ethanol for 2 min/step and Xylol for 1 min. Sections were air dried, mounted with Roti Histokitt (Roth) and covered with coverslips.

### Immunostaining

Free floating sections were transferred to a new 24-well plate and washed 2 x 10 min with 1x PBS. Nonspecific binding was prevented by incubating the sections in blocking buffer (5% BSA, 0.3% Triton-X100 in 1x PBS) for 60 min at room temperature. Then, the sections were incubated with primary antibodies (352, 1:500; Lamp1, 1:200; Arl8b, 1:200; 6E10 Alexa 594, 1:200) diluted in blocking buffer, overnight at 4 °C with gentle agitation. Sections were washed 3 times for 10 min with 1x PBS and incubated for 1 h at room temperature with the Alexa-fluorophor conjugated secondary antibody (1:1,000) diluted in blocking buffer. Then, the sections were washed three times for 10 min with 1x PBS and stained with Hoechst (1:5,000, ThermoFisher Scientific) diluted in 1x PBS for 20 sec. Sections were washed shortly in 1x PBS followed by ddH_2_O and mounted onto glass slides (Superfrost-Plus) using fluorescence mounting medium (Dako Agilent). Since the 6E10 antibody is conjugated to Alexa Fluor 594 (Alexa Fluor® 594 anti-β-Amyloid, 1-16), an incubation with a secondary antibody was not necessary. After incubation with Alexa 594 Fluor 6E10, brain sections were washed three times for 10 min with 1x PBS, mounted on slides, air dried shortly and directly used for Thioflavine S staining.

### Thioflavin S staining

Air dried sections were washed with PBS and incubated in 100, 95 and 80% EtOH for 2 min/step. Then, the sections were incubated in 1% Thioflavin S (MerckMillipore) solution for 15 min and washed with 50% EtOH for 1 min. Sections were briefly washed by 2 to 3 times dipping into H_2_O and air dried shortly. One drop of fluorescence mounting medium (Dako Agilent) was added and sections were sealed with a coverslip (24×50 mm).

### Microscopic imaging and image processing

Whole hemi brain sections were imaged using a Leica SP8 confocal laser scanning microscope (Leica Microsystems GmbH, Wetzlar). For quantification of plaques Thioflavin S dye was excited with the 405 nm diode laser and detected at 520 to 550 nm and Alexa Fluor 594-conjugated antibodies were excited with a 561 nm diode laser and detected at 610 to 700 nm. For correlation of Arl8b and 6E10 volumes, Alexa Fluor 594-conjugated antibodies were excited with a 561 nm laser and detected at 566 to 613 nm and Alexa Fluor 647 secondary antibodies staining Arl8b was excited with a 633 nm diode laser and detected at 640 to 700 nm. For detection, photomultiplier tubes (PMT) and Z-stack imaging was used. For quantification of plaques, eight sections per microscope slide were imaged in one workflow; images were acquired with a Plan Apo 20x/0.75 NA dry objective and the following settings: scan format 512 x 512 pixels, zoom 1x, pixel size of 1,137 µm and a z-step of 0,685 µm, scanner frequency of 600 Hz, pinhole at 1 Airy Unit and bi-directional scan to increase speed. All images were acquired with identical settings.

To achieve data sets with higher resolution (**Fig. 1f**) images where acquired with a Plan Apo 63x/1.4 NA Oil Objective, a scan format of 2048 x 2048 pixels, scanner frequency of 100 Hz, line average of 2 and pinhole at 1 Airy Unit. The image sampling parameter where adjusted to the Nyquist-Shannon theorem (pixel size of 0.05 µm and z-step of 0.15 µm, zoom of 1.8). To improve signal to noise ratio for stitching of tiles, image restoration (deconvolution by CMLE algorithm) was done subsequent to imaging by using Huygens Professional Suite (Scientific Volume Imaging, SVI).

**Figure 1.**
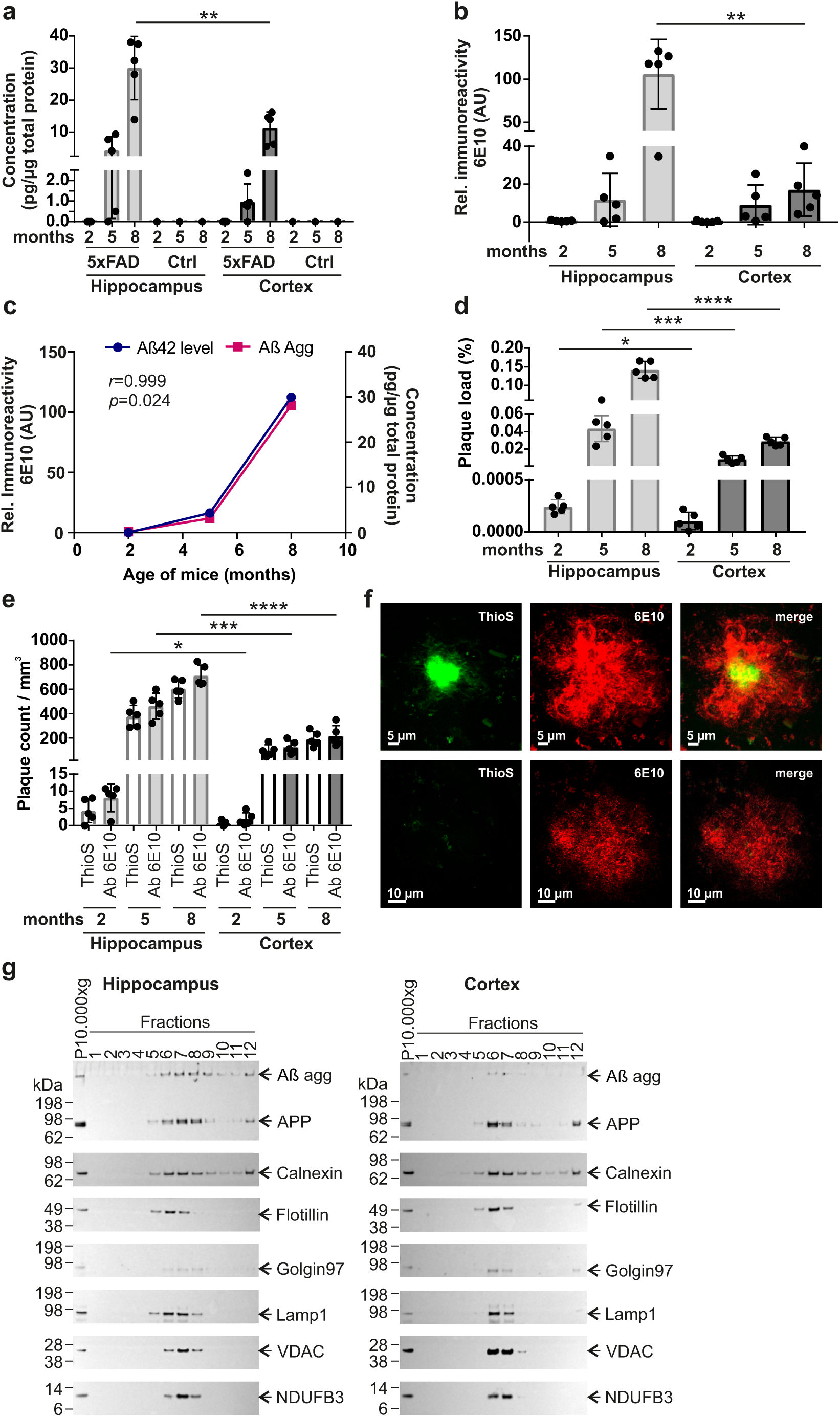
Analysis of Aβ -peptide levels and -aggregation in brains of 5xFAD mice. **(a)** Hippocampal and cortical brain extracts of 2-, 5- and 8-month-old AD (n=5) mice were used for Aβ42 peptide ELISAs. As a control (Ctrl), a pool of wt mouse brain extracts was tested in parallel, which was produced by mixing equal protein amounts of tissue extracts derived from 5 different wt mice per age and tissue. The graph represents the mean ± SD of five biological replicates of tg mice per age and tissue. Statistical analysis: Unpaired, two-tailed t test between hippocampal and cortical samples of mice with the same age (**, p = 0.0054). **(b)** Quantification of Aβ aggregates retained on filter membranes (**Additional File 1: Fig. S2a**) was performed using the Aida image analysis software. Per tissue and age, extracts of five different mice were analyzed. All data are expressed as mean ± SD of five biological replicates. Statistical significance was assessed between hippocampal and cortical samples of mice with the same age using an unpaired, two-tailed t test (**, p = 0.0016). **(c)** Pearson correlation between Aβ42 peptide levels determined by ELISA (blue line, right axis) and Aβ aggregates determined by MFA (purple line, left axis); hippocampal tissue samples were analyzed. The statistical significance of the association between the Aβ42 peptide levels and Aβ aggregates was measured with a two-tailed t-test (*, p = 0.024). **(d)** Plaque load was determined in hippocampus and cortex of 5xFAD mice at different ages (n = 5 mice per age and tissue). Plaques were immunohistochemically stained with the antibody 6E10. Data represent mean ± SD. The statistical significance was assessed between hippocampal and cortical tissues from the same age using an unpaired, two-tailed t test (*, p = 0.0204; ***, p = 0.0009; ****, p < 0.0001). **(e)** Plaque counts were determined in hippocampal and cortical tissues of 2-, 5- and 8-month-old 5xFAD mice (n=5 per age and tissue) by antibody 6E10 and ThioS staining. Data represent mean ± SD. The statistical significance was assessed with the antibody 6E10 (*, p = 0.0141; ***, p = 0.0002; ****, p < 0.0001) and ThioS staining (ns; ***, p = 0.0003; ****, p < 0.0001) each for hippocampal and cortical tissues of mice with the same age using an unpaired, two-tailed t test. **(f)** Immunofluorescence analysis of dense-core (top row) and diffuse plaques (bottom row) in a brain slice of an 8 month-old 5xFAD mouse. Red indicates 6E10 immunoreactive material. Green indicates fibrillar Aβ material stained with ThioS. Yellow indicates merged signal. **(g)** Fractionation of hippocampal and cortical brain extracts derived from 8-month-old 5xFAD mice. Fractions from sucrose density gradient centrifugations of 10.000xg membrane pellets were analysed by immunoblotting using antibodies detecting APP and amyloid-β (antibody 6E10), marker proteins of the endoplasmic reticulum (Calnexin), lipid rafts (Flotillin), the Golgi apparatus (Golgin97), lysosomes (Lamp1) and mitochondria (VDAC, NDUFB3). Equal exposure times per antibody for hippocampus and cortex are shown. Per fraction, equal volumes were loaded. From the solubilized pellet (P10.000xg) 5 µg was loaded.

### Plaque quantification

Plaque counts and plaque volumes were quantified using Imaris 9.2.1 surface function for surface detection. The detection threshold for each channel (6E10 or ThioS) was set by hand and slightly adjusted within a range of a reference point to obtain values as constant as possible. Plaques smaller than 50 µm^3^ were excluded from quantification. The total area of hippocampus and cortex was measured in each section and normalized with 40 µm section thickness to determine their respective volume. Artefacts from staining, sectioning or handling were excluded manually.

Plaques were separately quantified per section in each region and the respective plaque volumes and plaque counts were determined. Based on these, the plaque load was calculated as the proportion of the hippocampal or cortical volume occupied by plaques: Aβ Plaque Load (%) = (Total Volume Plaques/Total Volume Region) × 100. For quantification of Arl8b volume, Imaris 9.6 Software (Bitplane) was used. The detection threshold was set manually and slightly adjusted within the range of a reference point to maintain values as constant as possible.

### Reverse transcription und quantitative real-time PCR

To purify RNA from hippocampus and cortex the RNeasy Lipid Tissue kit (Qiagen) was used. DNA contamination was removed from purified RNA by using the RNase-Free DNase set (Qiagen). RNA concentration was determined by measuring the absorbance at 260 nm. Single stranded cDNA from total RNA was synthesized using the High Capacity cDNA Reverse Transcription kit (ThermoFisher Scientific). For Real-Time PCR of human APP, 10 ng cDNA was amplified with a TaqMan gene expression assay (ThermoFisher Scientific: Hs00169098_m1) and TaqMan Gene Expression Master Mix (ThermoFisher Scientific) in a total volume of 10 µl. For normalization of data, an endogenous mouse EIF-4H (Mm00504282_m1) FAM/MGB labelled probe (ThermoFisher Scientific) was used. For real-time PCR of human *PSEN1*, 10 ng cDNA was amplified with a TaqMan gene expression assay (ThermoFisher Scientific: Hs00997789_m1) and TaqMan Gene Expression Master Mix (ThermoFisher Scientific) in a total volume of 10 µl. For normalization of data, an endogenous mouse *GAPDH* (Mm99999915_g1) FAM/MGB labelled probe (ThermoFisher Scientific) was used. Real-time PCR was performed using the ViiA 7 real-time PCR system (Applied Biosystems). Samples were measured in triplicates. Quantification was performed using the ΔCt method and the QuantStudio real-time PCR Software v1.3.

### Preparation of mouse brain samples for mass spectrometric analysis

Mouse brain material was weighed and homogenized in lysis buffer (6 M guanidinium chloride, 100 mM Tris pH 8.5, 10 mM TCEP, 40 mM CAA, 20% weight/volume), heated for 5 min at 95 °C, cooled on ice for 15 min, followed by sonication. After centrifugation for 30 min at 3,500g (4 °C), the supernatant (soluble protein fraction) was transferred to a fresh tube and mixed with 4 volumes of ice cold acetone, followed by incubation over night at -20 °C. Samples were centrifuged for 15 min at 12,000g (4 °C) and the resulting protein pellets were washed with ice cold 80% acetone, air dried and resuspended in digestion buffer (6 M urea, 2 M thiourea, 100 mM Hepes, pH 8). The samples were sonicated using a Branson probe Sonifier^TM^ (output 3-4, 50% duty cycle, 4× 30 s). Protein concentration was determined using a Bradford assay (BioRad). Samples were stored at -80 °C until use. 50 μg protein per sample were used for tryptic digestion. First, Endopeptidase LysC (Wako, Japan) was added in a protein:enzyme ratio of 50:1 and incubated for 4 hours at room temperature. After dilution of the sample with 4 volumes 50 mM ammonium bicarbonate (pH 8.0), sequencing grade modified trypsin (Promega) was added (protein:enzyme ratio 100:1) and digested overnight at room temperature. Trypsin and Lys-C activity was quenched by acidification with TFA to pH ∼2. Peptides were cleaned up using the StageTip protocol [38].

### Mass spectrometric analyses

For mass spectrometric analyses peptide samples (2 µg per measurement) were separated by reversed phase chromatography using the Eksigent NanoLC 400 system (Sciex) on in-house manufactured 20 cm fritless silica microcolumns with an inner diameter of 75 µm, packed with 3 µm ReproSil-Pur C18-AQ beads. A 8-60% acetonitrile gradient (224 min) at a nanoflow rate of 250 nl/min was applied. Eluting peptides were directly ionized by electrospray ionization and analyzed on a Thermo Orbitrap Fusion (Q-OT-qIT, Thermo). Survey scans of peptide precursors from 300 to 1,500 m/z were performed at 120K resolution with a 2 × 10^5^ ion count target. Tandem MS was performed by isolation at 1.6 m/z with the quadrupole, HCD fragmentation with normalized collision energy of 30, and rapid scan MS analysis in the ion trap. The MS^2^ ion count target was set to 2×10^3^ and the maximum injection time was 300 ms. Only precursors with charge state 2–7 were sampled for MS^2^. The dynamic exclusion duration was set to 60 s with a 10 ppm tolerance around the selected precursor and its isotopes. The instrument was run in top speed mode with 3 second cycles, meaning the instrument would continuously perform MS^2^ events until the list of nonexcluded precursors diminishes to zero or 3 s. For all samples, 2 technical replicates were performed (for 5 months hippocampus, 3 technical replicates were measured).

### Analysis of mass spectrometric data

Data were analyzed by MaxQuant software (v1.5.1.2). The internal Andromeda search engine was used to search MS^2^ spectra against a decoy human UniProt database (MOUSE.2014-10) containing forward and reverse sequences. The search included variable modifications of methionine oxidation and N-terminal acetylation, deamidation (NQ) and fixed modification of carbamidomethyl cysteine. Minimal peptide length was set to seven amino acids and a maximum of two missed cleavages was allowed. The FDR was set to 1% for peptide and protein identifications. The integrated LFQ quantitation algorithm was applied. Unique and razor peptides were considered for quantification with a minimum ratio count of 1. Retention times were recalibrated based on the built-in nonlinear time-rescaling algorithm. MS^2^ identifications were transferred between runs with the “Match between runs” option, in which the maximal retention time window was set to 2 min.

### Quantification of differential protein abundance

The LFQ intensities obtained from MaxQuant were batch-corrected separately for cortex and hippocampus data using the Combat [39] algorithm on log10 LFQ intensities. No missing intensity values imputation was performed. Instead, we adopted a conservative approach, keeping for analysis only proteins with reliable intensity measures across the majority of conditions. In a first selection step, we grouped the data by tissue, age and genotype. When a protein had missing intensity values in half of the samples or more for each individual condition, it was deemed unreliable and was excluded. The number of proteins reliably measured in the cortex ranged from 3602 to 3752 (median 3651), as 4915 proteins were quantified in this dataset, i.e. 1163 and 1313 (median 1264) proteins were discarded. In the hippocampus dataset 5069 proteins were quantified and between 3434 and 3620 (median 3502) proteins were deemed reliably measured according to the described threshold. In the next step, we ensured that protein intensities were reliably measured in both 5xFAD and control samples collected in the same tissue and at the same age. The number of proteins reliably measured in both genotypes are 3613 (2 months), 3476 (5 months) & 3524 (8 months) in cortex, and 3337 (2 months), 3471 (5 months) and 3248 (8 months) in hippocampus. These proteins were used for the analysis of the transgene effect, when it was restricted to samples collected at the same age and in the same tissue. In total, we identified 699 differentially expressed proteins defined through the “pairwise model” (**Fig. 2b, Additional File 3: Supplementary Excel File 1**) that are significantly dysregulated at 2, 5 or 8 months in at least one tissue. DEPs across all time points (DE.cortex.Age2, DE.cortex.Age5 and DE.cortex.Age8; DE.hippo.Age2, DE.hippo.Age5 and DE.hippo.Age8) are presented as volcano plots (**Fig. 2a**) that were generated with the ggplot2 package in R v.4.2.0. The volcano plots show the protein expression logarithmic fold-changes (log10 FC; x-axis) and the adjusted p-values (-log10 p-value; y-axis). However, these pairwise models do not allow comparisons of the transgene effect across ages or tissues.

**Figure 2.**
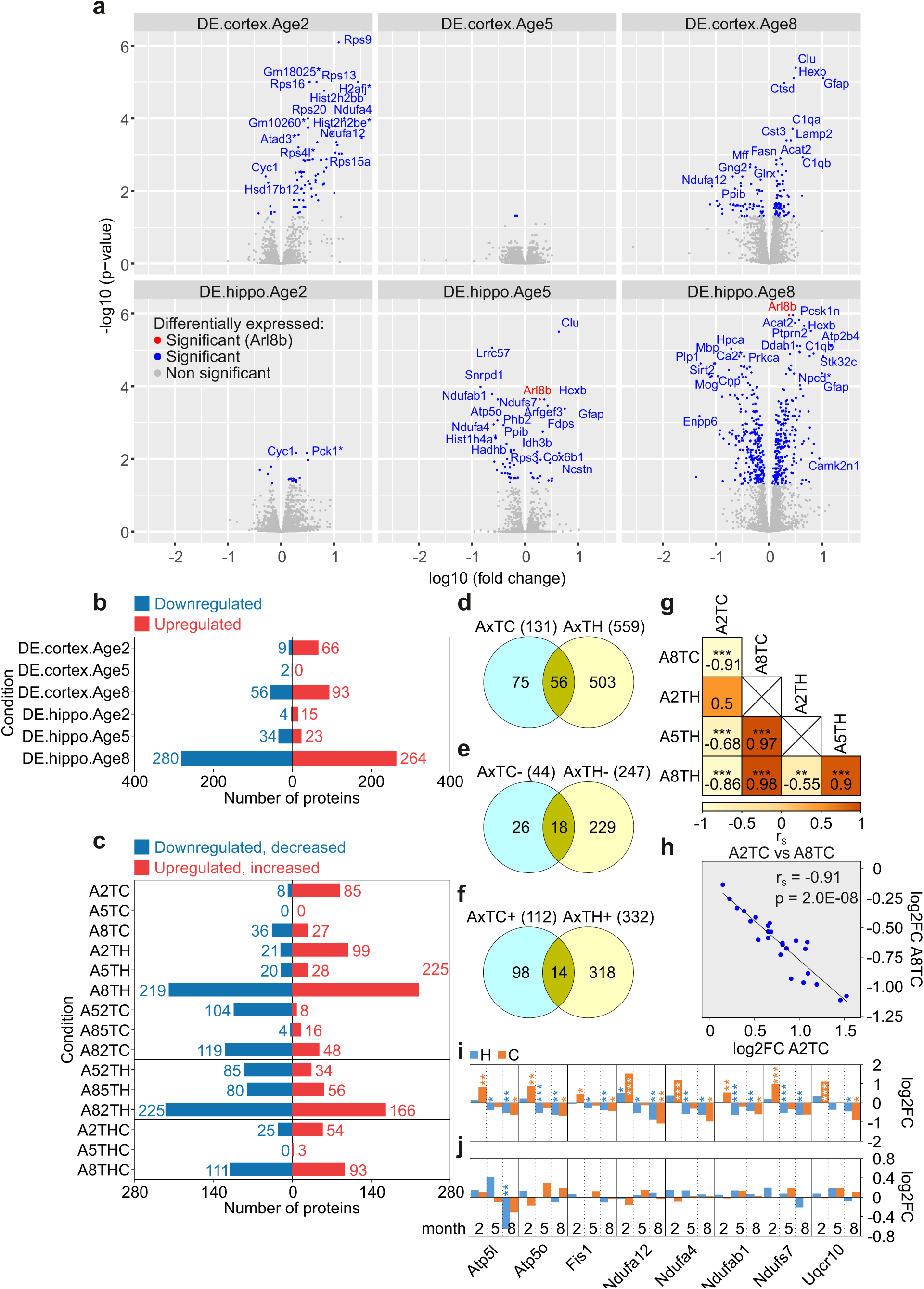
Differential expression analysis of proteins in brains of 5xFAD mice. **(a)** Volcano plots depicting the protein expression logarithmic fold-changes (log10 fold change, x-axis) and the adjusted p-values (-log10 p-value, y-axis). **D**ifferentially **E**xpressed proteins across all time points (**Age 2**, **5** and **8**) in **cortex** and **hippo**campus were analyzed (DE.cortex.Age2, DE.cortex.Age5 and DE.cortex.Age8; DE.hippo.Age2, DE.hippo.Age5 and DE.hippo.Age8). All proteins highlighted in blue have significantly altered expression. The candidate AD biomarker Arl8b, marked in red, is significantly upregulated in hippocampus. Gene names for potential proteins of interest with highly significant fold changes are indicated. Proteins not significantly changed in their abundance are highlighted in grey. Significance was determined using a two-tailed t test and a Benjamini-Hochberg False Discovery Rate (FDR) set to 5%. **(b)** Numbers of differentially expressed proteins obtained by comparing the protein expression measurements of 5xFAD mice at different ages with their corresponding, age-matched, wild-type controls in cortex and hippocampus. The identifiers of the amounts of proteins analyzed were denoted analogously as in panel (a). Bars indicate the numbers of significant down-(blue) and up-regulated (red) proteins. **(c)** Numbers of proteins significantly differentially regulated. A2TC (**A**ge **2**, **T**issue **C**ortex), A5TC and A8TC label the numbers of proteins for which the transgene effect is significant in cortex, at 2, 5 and 8 months, respectively. Proteins more abundant in 5xFAD mice are shown in red, while proteins more abundant in wild-type mice are shown in blue. A2TH (**A**ge **2**, **T**issue **H**ippocampus), A5TH and A8TH label the same comparison in hippocampus. A52TC (**A**ge **5** vs **2**, **T**issue **C**ortex), A85TC and A82TC refer to the numbers of proteins for which the transgene effect is significantly different at two time points (5 vs 2, 8 vs 5 and 8 vs 2 months) in cortex. The numbers of proteins which increase in abundance with age are shown in red, while the numbers of proteins that decrease with age are shown in blue. The corresponding labelling for the hippocampus samples are A52TH (**A**ge 5 vs **2**, **T**issue **H**ippocampus), A85TH and A82TH. Finally, A2THC (**A**ge **2**, **T**issue **H**ippocampus vs **C**ortex), A5THC and A8THC report the numbers of proteins for which the transgene effect is significantly different in cortex and hippocampus at 2, 5 and 8 months, respectively. Higher hippocampus abundance is shown in red on the right, and higher cortex abundance in blue on the left. The underlying data are available as Additional File 3: Supplementary Excel File 2. **(d-f)** Venn diagrams showing the numbers of total **(d)**, downregulated **(e)** and upregulated **(f)** proteins in cortex and hippocampus, including only significantly differentially expressed proteins (DEPs). The amounts of DEPs were denoted analogously to panel (c) but were combined across all time points (months 2, 5 and 8); the combination of DEPs is indicated by the x (e.g., AxTC). **(g)** Correlation analysis of DEPs for A2TC, A8TC, A2TH, A5TH and A8TH. The degree of correlation was assessed by Spearman correlation coefficients (r_S_) and corresponding FDR adjusted p values (**, p < 0.01; ***, p < 0.001). Crosses indicate no correlation. The colors denote the values of the Spearman correlation coefficients. **(h)** Example Spearman inverse correlation of A2TC versus A8TC also shown in panel (g). **(i)** Time-dependent changes in the abundance of 8 mitochondrial transcripts (below) and proteins (top) that play a key role in oxidative phosphorylation and ATP production. The temporal changes of the LFCs across all ages (2, 5 and 8 months) in cortex (orange) and hippocampus (blue) are shown. The statistical significance of the differentially expressed proteins and transcripts was measured with a two-tailed t-test, adjusted by the Benjamini-Hochberg multiple testing correction (*, p<0.05, **, p < 0.01; ***, p < 0.001). All analyzes are based on mean values of measured intensities from five biological replicates of tg mice per age and tissue (n=5).

For this reason, we have created a set of proteins that are reliably measured in all conditions, suitable for all comparisons of transgene effects. For this set, we demanded that the proteins identified in the previous step should be reliably measured in each tissue, in all time points. These requirements reduced the numbers to 3281 and 3047 proteins for the cortex and hippocampus, respectively. Finally, only proteins common to the filtered cortex and hippocampus datasets were included in the statistical model. Please note that the Andromeda assignment of peptides to proteins may differ between cortex and hippocampus datasets. As a result, some peptides are attributed to several proteins and in rare cases this may lead to ambiguous mapping of proteins. Therefore, we discarded any many-to-many relationships between cortex and hippocampus proteins. Eventually, 2761 reliably measured proteins common to both tissues were used in the final full statistical model for differential protein analysis. While our filtering discards many proteins it allows the statistical estimation of transgene effect differences between conditions. In total, we identified 637 differentially expressed proteins defined through the “full model” (**Fig. 2c, Additional File 3: Supplementary Excel File 2**) that are significantly dysregulated at 2, 5 or 8 months in at least one tissue (A2TC, A5TC or A8TC; A2TH, A5TH or A8TH).

To estimate the transgene effect, we analysed differential intensities on log10 scale. Differential protein abundance between wild-type and 5xFAD animals are estimated separately at each age group and in each tissue, using empirical Bayes statistics implemented in limma [40]. Replicated measures on the same animal were taken into account using the ‘duplicateCorrelation’ procedure in limma. The differences in transgene effect between tissues or time points were obtained using contrasts to take the interaction effect into account. The Benjamini-Hochberg False Discovery Rate [41] was computed separately for each contrast. Proteins were considered differentially expressed when their FDR was below 0.05. To make sure that the aggregation of data of very different conditions does not distort the results obtained from the model, we compared transgene effects obtained using the full model to those obtained from pairwise models in each condition separately. Pearson correlation between *t* statistics estimated from the full model and the pairwise models are between 0.942 and 0.977 (median 0.955). This suggests that including data from different tissues does not have a strong influence on the transgene effect modelling.

### Generation of Venn diagrams showing the logical relation between datasets of differentially expressed proteins

The Venn diagrams (**Fig. 2d-f, 3e, 4a**) were generated with Venny 2.0 (https://bioinfogp.cnb.csic.es/tools/venny/index2.0.2.html) and Corel Draw. In figures 2d-f they show the numbers of total, downregulated and upregulated DEPs in cortex and hippocampus defined through the “full model” (**Additional File 3: Supplementary Excel File 2**). The amounts of DEPs were combined across all time points (months 2, 5 and 8) and are indicated by the x (e.g., AxTC). Figure 3e represents the number of differentially expressed genes (DEGs, **Additional File 3: Supplementary Excel File 4**) that overlap with Aβ aggregate-correlated and anti-correlated (acorr) DEPs (**Additional File 3: Supplementary Excel File 3**). In figure 4a the overlaps of differentially expressed proteins in brains of 5xFAD mice from the “pairwise model” (**Additional File 3: Supplementary Excel File 1**) and post-mortem brains of AD patients from Drummond 2022 ([39], Table S1), Johnson 2020 ([40], Supplementary Table 2A) and Johnson 2022 ([41], Supplementary Table 2) were investigated.

**Figure 3.**
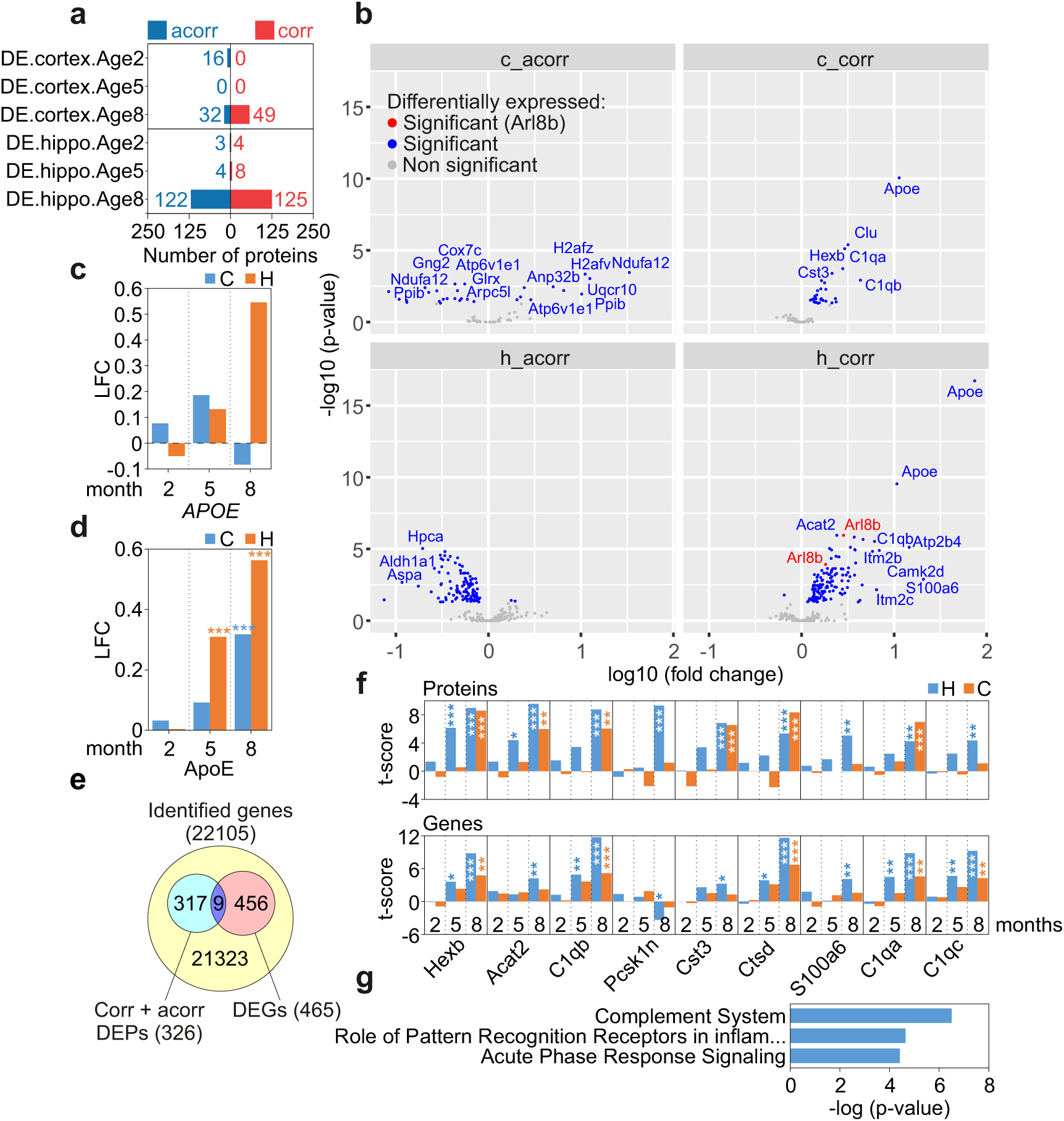
Identification of Aβ correlated and anticorrelated protein alterations in 5xFAD brains. **(a)** Numbers of proteins that correlate (corr, in red) or anticorrelate (acorr, in blue) with Aβ aggregates in hippocampus and cortex and are differentially expressed at 2 (DE.cortex.Age2, DE.hippo.Age2), 5 (DE.cortex.Age5, DE.hippo.Age5) or 8 (DE.cortex.Age8, DE.hippo.Age8) months. The identifiers were denoted analogously as in figure 2a. The degree of correlation was assessed by Pearson correlation and corresponding FDR adjusted p values. **(b)** Volcano plots depicting the protein expression logarithmic fold-changes (log10 fold change, x-axis) and the adjusted p-values (-log10 p-value, y-axis) across all time points (age 2, 5 and 8 months) for DEPs that correlate (corr) or anticorrelate (acorr) with Aβ aggregates in hippocampus (h) and cortex (c). All proteins highlighted in blue are expressed significantly differently. The Aβ correlating protein AD biomarker candiate Arl8b is marked in red. Proteins of interest with highly significant fold changes are indicated with gene names. Proteins not significantly changed are highlighted in grey. Significance was determined using a two-tailed t test and a Benjamini-Hochberg False Discovery Rate (FDR) set to 5%. **(c, d)** Changes of *APOE* transcript **(c)** and protein **(d)** levels in hippocampus (H) and cortex (C) of 2-, 5-, and 8-month-old 5xFAD mice. LFC, log2 fold change. **(e)** Number of differentially expressed genes (DEGs, light red) that overlap (purple) with Aβ aggregate-correlated (corr) and anticorrelated (acorr) DEPs (light blue). **(f)** Time-dependent changes in the abundance of 9 correlating or anticorrelating molecules; both protein (top) and transcript (bottom) level changes are shown. The temporal changes of the t-scores across all ages (2, 5 and 8 months) in cortex (orange) and hippocampus (blue) are illustrated. The statistical significance of differentially expressed proteins and transcripts was measured with a two-tailed t-test, adjusted by the Benjamini-Hochberg multiple testing correction (*, p<0.05, **, p < 0.01; ***, p < 0.001). **(g)** Ingenuity pathway analysis (IPA) for the correlating or anticorrelating molecules that are changed both at the protein and transcript level as shown in panels (e) and (f). The statistical significance of the association between the DEPs and the canonical pathways was measured with a right-tailed Fisher’s exact test to calculate the p-values, adjusted by the Benjamini-Hochberg multiple testing correction. All analyzes are based on mean values of measured intensities from five biological replicates of tg mice per age and tissue (n=5).

**Figure 4.**
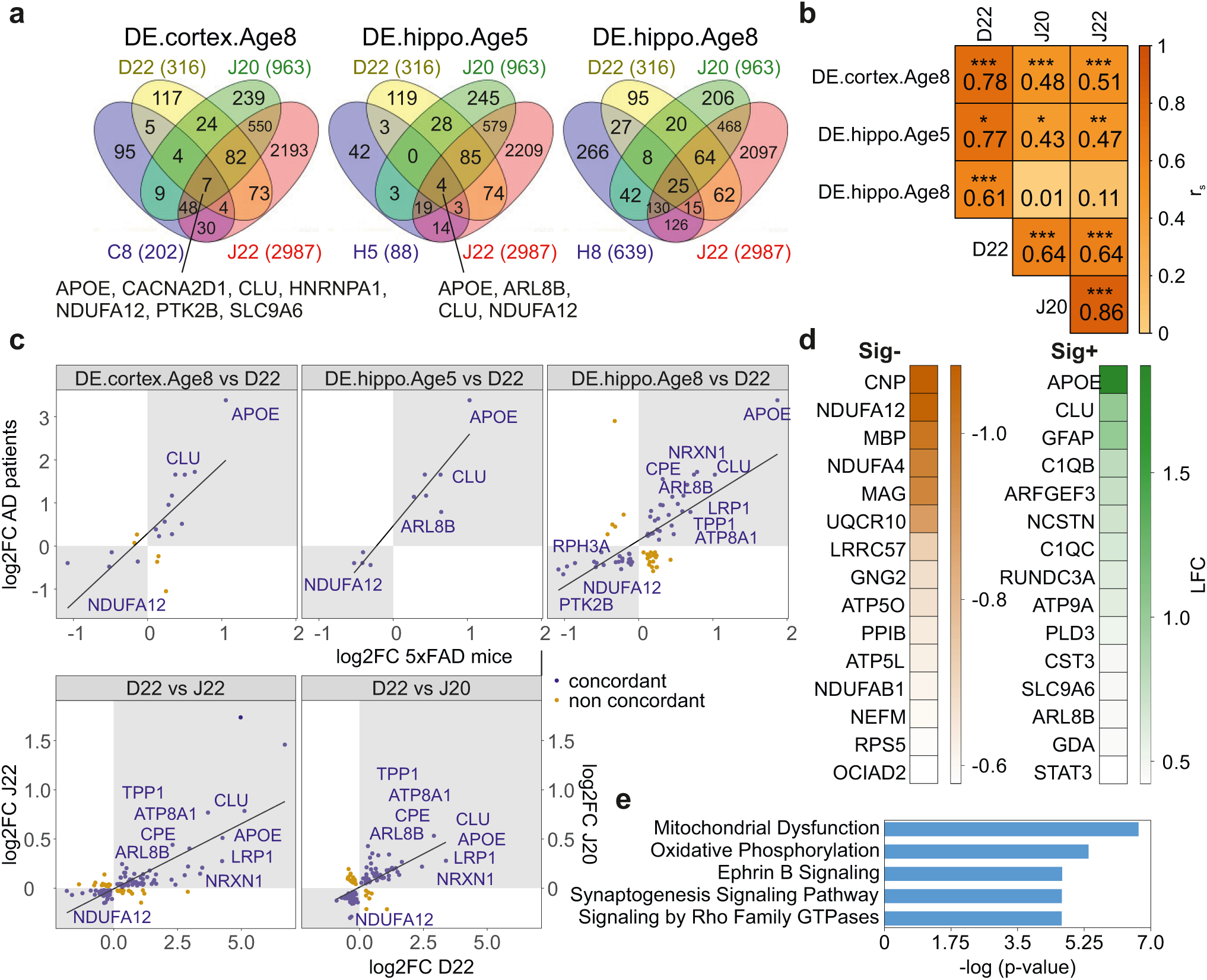
Analysis of dysregulated proteins of 5xFAD brains and postmortem brains of AD patients. **(a)** Investigation of the overlap of DEPs in brains of 5xFAD mice and postmortem AD patients. Protein changes in cortical and hippocampal tissues of 8- and 5-month-old mice (DE.cortex.Age8; DE.hippo.Age5; DE.hippo.Age8) that contain Aβ aggregates (Fig. 1d) were compared with patient protein measurements. The amounts of DEPs are denoted analogously to figure 2a or have been abbreviated (C8, H5, H8). The human AD data were obtained from Drummond 2022 (D22, [65]), Johnson 2020 (J20, [64]) and Johnson 2022 (J22, [63]). The total numbers of DEPs in each dataset are indicated in brackets. In DE.cortex.Age8 and DE.hippo.Age5 the proteins present in all four data sets are depicted with gene names. **(b)** Correlation analyses of DEPs from cortical tissues of 8-month-old mice (DE.cortex.Age8) and DEPs from hippocampal tissues of 5-(DE.hippo.Age5) and 8-month-old mice (DE.hippo.Age8) with human AD patient data from Drummond 2022, Johnson 2020 and Johnson 2022 (as indicated in panel (a)) were performed. The degree of correlation was assessed by Spearman correlation coefficients (r_S_) and corresponding FDR adjusted p values (*, p<0.05; **, p < 0.01; ***, p < 0.001). Colours denote the values of the Spearman correlation coefficients. **(c)** Example Spearman correlations and concordance representations of dysregulated proteins from 5xFAD mice (DE.cortex.Age8, DE.cortex.Age5 and DE.cortex.Age8) versus human AD patient data from Drummond 2022 along with human AD data against each other (D22 vs J22, D22 vs J20) are shown. The identifiers are denoted analogously to figures 2a and 4a. Proteins concordantly up- or downregulated in 5xFAD mice and human AD patient brains are shown in grey quadrants and marked in blue. Proteins of interest with highly significant fold changes are indicated with gene names. Correlating but non-concordant proteins are marked in brown. **(d)** Generation of signatures for proteins concordantly dysregulated in 5xFAD mice and human AD patient brains. The heatmaps show the 15 most down (Sig-, left) or upregulated (Sig+, right) proteins. Colors denote the values of the log2 fold changes. **(e)** Ingenuity pathway analysis (IPA) for the complete set of concordantly dysregulated proteins. The statistical significance of the association between the DEPs and the canonical pathways was measured with a right-tailed Fisher’s exact test to calculate p-values, adjusted by the Benjamini-Hochberg multiple testing correction. All analyzes are based on mean values of measured intensities from five biological replicates of tg mice per age and tissue (n=5).

### RNA extraction and sequencing

The brain tissue was homogenized in TRIzol® Reagent (Ambion) using a Precellys homogenizer (Bertin Technologies) and extracted according to the TRIzol® Reagent protocol, with an additional phenol/chloroform purification step and the final isopropanol precipitation. For the total RNA sequencing we used 1,000 ng of total RNA; rRNA was depleted using a RNase H-based protocol. We mixed total RNA with 1 μg of a DNA oligonucleotide pool comprising a 50-nt long oligonucleotide mix covering the reverse complement of the entire length of each human rRNA (28S rRNA, 18S rRNA, 16S rRNA, 5.8S rRNA, 5S rRNA, 12S rRNA), incubated with 1U of RNase H (Hybridase Thermostable RNase H, Epicentre), purified using RNA Cleanup XP beads (Agencourt), DNase treated using TURBO DNase rigorous treatment protocol (Thermo Fisher Scientific), and purified again with RNA Cleanup XP beads. We fragmented the rRNA-depleted RNA samples and processed them into strand-specific cDNA libraries using TruSeq Stranded Total LT Sample Prep Kit (Illumina) and then sequenced them on NextSeq 500, High Output Kit, 1 x 150 cycles.

### Quantification of differential gene abundance

RNA-Seq reads were mapped to the mouse GRCm38.p4 genome with STAR [39] (version 2.4.2.a). Reads were assigned to genes with featureCounts (version 1.4.6-p5) using the following parameters: -t exon -g gene_id, and with Genecode grcm38.p4-vM6 gtf reference [40]. The differential expression analysis was carried out with DESeq2 [41] (version 1.12.4) using default parameters.

### Mapping of gene identifiers

The joint analysis of proteomics and transcriptomics data was done on genes. The LFQ intensity, the differential protein abundance and associated statistical quantities were attributed to a gene by the following process: first, each protein identifier produced by the MaxQuant analysis software was assigned to a MGI gene identifier. This was done by matching the MaxQuant protein identifier with GenBank, UniProt and TrEMBL protein ids obtained from MGI makers association files [42] downloaded in August 2017. This mapping provided connections between the initial protein identifier to NCBI & ENSEMBL gene identifiers. For protein ids unknown in MGD, direct queries to the UniProt database were made, and ENSEMBL biomart [43] was used to link the UniProt output to ENSEMBL & NCBI gene ids.

### Correlation analysis methods

To evaluate the relationship between the abundance changes of the differentially expressed proteins (DEPs) and the progressive Aβ aggregate formation in cortex and hippocampus of 5xFAD brains, we used the Pearson correlation coefficient r. We examined how the abundance changes of all 699 DEPs (**Fig. 2a, b, Additional File 3: Supplementary Excel File 1**) are correlated or anticorrelated with the time-dependent accumulation of Aβ aggregates by associating the LFCs of the measured intensities of the DEPs against the abundance of Aβ aggregates across all ages (months 2, 5 and 8). Four different Aβ datasets were used: amounts of Aβ aggregates determined by MFA (**Fig. 1b**), Aβ plaque load (**Fig. 1d**), Aβ plaque counts determined by 6E10 antibody and ThioS staining (**Fig. 1e**). Each dataset consists of three data points, which are mean values of measured intensities from 5 biological replicates per age. For the correlation analysis, we examined the relationship between the three data points of the DEPs and the Aβ data sets separately for cortex and hippocampus. As the result, only DEPs with changes that correlate (0 < r < 1) or anticorrelate (0 > r > -1) significantly (p_corr_ < 0.05) with at least one of the mentioned Aβ datasets were considered (**Fig. 3a, Additional File 3: Supplementary Excel File 3**). The results were summarized across all ages in four datasets: c_corr -Aβ-correlating proteins in cortex; c_acorr - Aβ-anticorrelating proteins in cortex; h_corr - Aβ-correlating proteins in hippocampus; h_acorr - Aβ-anticorrelating proteins in hippocampus (**Fig. 3b, Additional File 3: Supplementary Excel File 3**). The differential expression of the correlating and anticorrelating proteins (c_corr, c_acorr, h_corr, h_acorr) is presented in volcano plots (**Fig. 3b, Additional File 3: Supplementary Excel File 3**) that were generated with the ggplot2 package in R v.4.2.0. The volcano plots show the protein expression logarithmic fold-changes (log10 FC; x-axis) and the adjusted p-values (-log10 p-value; y-axis). Pearson correlation coefficients between the volumes of accumulated protein Arl8b and 6E10-stained amyloid beta plaques in hippocampus of 2-(H2), 5-(H5) and 8-(H8) month-old 5xFAD mice were computed with GraphPad Prism v7 (**Fig. 5f**). Temporal Pearson correlations between Aβ42 peptide levels determined by ELISA and Aβ aggregates detected by MFA in hippocampus (**Fig. 1c**) and cortex (**Additional File 2: Fig. S2b**) were calculated in GraphPad Prism v7. To analyse the association of CSF biomarkers [Aβ(1-42), Aβ(1-40), Aβ42/Aβ40, t-tau and p-tau] and CSF Arl8b concentrations a Spearman correlation using GraphPad Prism v7 was performed (**Additional File 1: Table S3**). For the correlation analysis Arl8b and biomarker values determined in CSF samples derived from AD patients and control individuals were used. The statistical significance of the association was measured with a two-tailed t-test.

**Figure 5.**
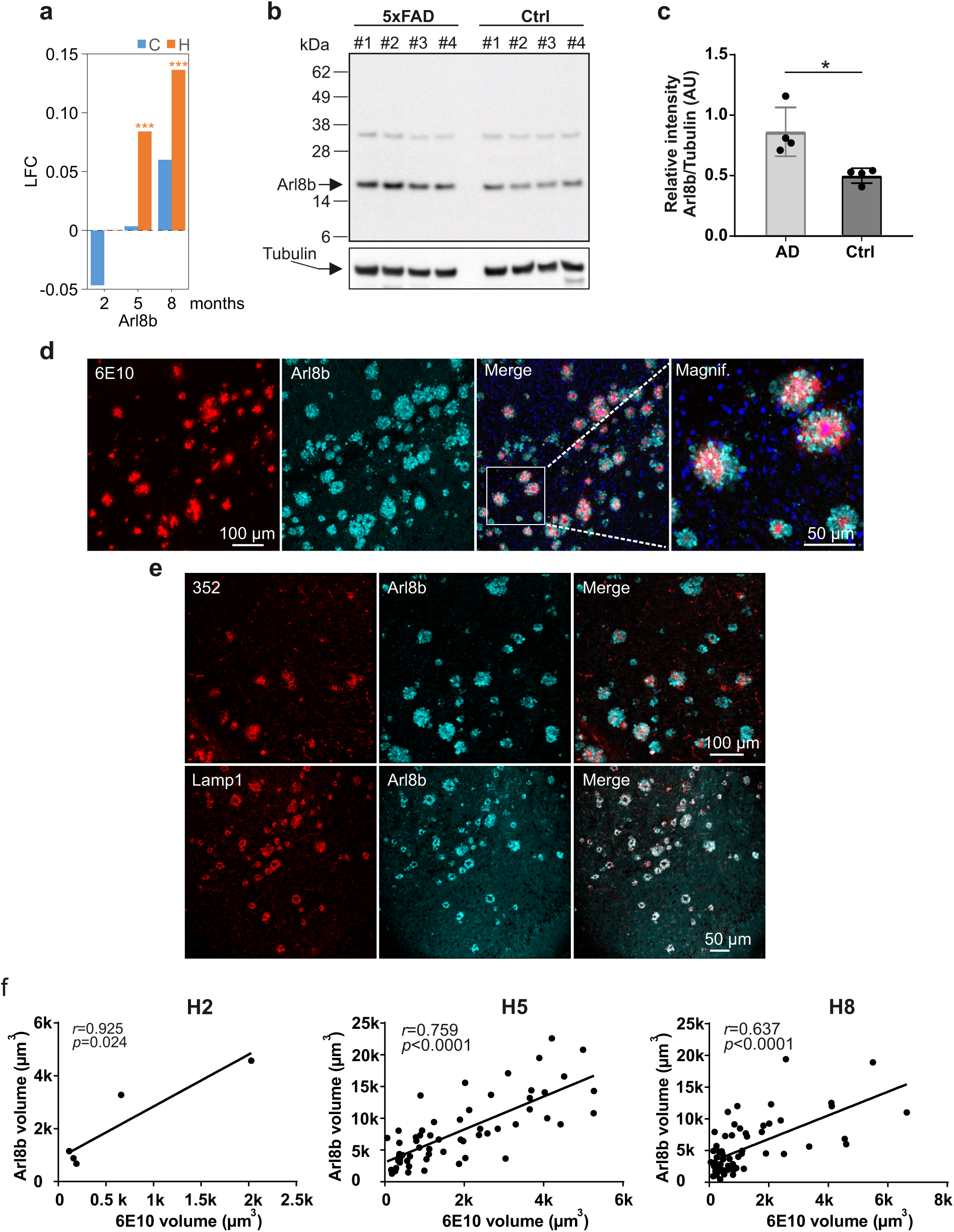
The protein Arl8b is upregulated in hippocampal tissues of 5xFAD mice. **(a)** Time-depended change of Arl8b protein abundance in hippocampal and cortical tissues of 5xFAD transgenic animals. LFC, log2 fold change. **(b)** Hippocampal brain homogenates prepared from four 8-month-old 5xFAD (1 to 4) and control (ctrl; 1 to 4) mice were analyzed by SDS-PAGE and immunoblotting using anti-Arl8b. As a control, a tubulin (anti-alpha Tubulin, #T6074) immunoblot was performed. **(c)** Quantification of Arl8b expression in relation to tubulin using band intensities of immunoblots in (**b**). Relative intensity values (mean ± SD) are shown for AD (n=4) and Ctrl (n=4) mice. Statistical significance was assessed between AD and Ctrl mice using an unpaired, two-tailed t test (*, p = 0.0137). **(d)** Immunofluorescence analysis of 5xFAD mouse (8 months) brain slices using AlexaFluor594-labelled 6E10 antibody (red); an anti-Arl8b antibody combined with an AlexaFluor647-labelled anti-rabbit IgG (turquoise) was applied to detect Arl8b. The scale bar shown in the 6E10 image also applies for the Arl8b and merge image. The picture on the right shows a magnification (magnif.) of an area indicated in the merged picture. **(e)** Brain slices of 8-month-old 5xFAD mice were stained with the primary antibodies indicated in the images. For detection with 352 and Lamp1 antibodies an AlexaFluor594-labelled anti-mouse IgG (red) was used; for Arl8b detection an AlexaFluor647-labelled anti-rabbit IgG (turquoise) was applied. Antibody 352 specifically recognizes Aβ42 fibrillar aggregates [30]. **(f)** Pearson correlation between the volumes of Arl8b accumulations and 6E10-stained amyloid beta plaques in hippocampus of 2-(H2), 5-(H5) and 8-(H8) month-old 5xFAD mice. A total of 60 plaques were analyzed in brain slices derived from 5- and 8-month-old 5xFAD mice. For 2-month-old 5xFAD mice less than 60 plaques were analyzed, since amyloid burden at this age is low. The statistical significance of the association between the volumes of Arl8b accumulations and 6E10-stained amyloid-b plaques was measured with a two-tailed t-test (*, p = 0.024; ****, p < 0.0001).

### Generation of correlation matrices of selected differentially expressed proteins

In order to compute the Spearman correlation between DEPs of selected single condition datasets (A2TC, A8TC, A2TH, A5TH and A8TH) in 5xFAD mice (**Additional File 3: Supplementary Excel File 2**) a correlation matrix was generated (**Fig. 2g**) using the packages correlation and corrplot in R v.4.2.0. Since no proteins are significantly changed in the cortex at 5 months (A5TC) this data set is omitted in the correlation analysis. To illustrate an example of strongly associated datasets from figure 2g, the correlation between the datasets A2TC and A8TC is shown as a scatterplot (**Fig. 2h**) representing the expression logarithmic fold-changes (log2 FC) of the DEPs in both datasets. The scatterplot was generated using the ggplot2 package in R v.4.2.0. In figure 4b the correlation analysis of the DEPs from cortical tissues of 8-month-old mice (DE.cortex.Age8) and DEPs from hippocampal tissues of 5-(DE.hippo.Age5) and 8-month-old mice (DE.hippo.Age8) with human AD patient data from Drummond 2022 ([39], Table S1), Johnson 2020 ([40], Supplementary Table 2A) and Johnson 2022 ([41], Supplementary Table 2) was performed. Strong correlations were selected from figure 4b in order to represent them qualitatively in figure 4c and to illustrate the concordance using the ggplot2 package in R v.4.2.0. The proteins that are concordantly up- or downregulated in 5xFAD mice and human AD patient brains are shown.

### Generation of protein signatures

The strategy to generate signatures for proteins that are concordantly dysregulated in 5xFAD mice and human AD patient brains is shown schematically in **Additional File 2: Fig. S8**. Heatmaps were created with Graphpad Prism v7 showing the 15 most down (Sig-, left) and upregulated (Sig+, right) proteins (**Fig. 4d, Additional File 3: Supplementary Excel File 6**). The top dysregulated proteins were selected from the complete set of proteins that are concordantly dysregulated (**Additional File 3: Supplementary Excel File 6**).

### Enrichment analysis of the dysregulated proteins in brains of 5xFAD mice for **specific cell types**

The proteomics data from Sharma et al. [44] were used to determine protein markers, expressed exclusively in a subset of brain cells (**Additional File 2: Fig. S5**). The markers are defined as genes with at least 10-fold higher expression in one cell type than in the average of the other cell types. Only the sets of differentially expressed proteins from the pairwise model were analysed for the enrichment of specific cell types.

### Differential centrifugation of mouse brain homogenates and sucrose density gradient centrifugation

The protocol was adopted from a previously reported study [45] with little modifications. Four hippocampi and 4 cortices from 8-month-old 5xFAD mice were homogenized in ice cold sucrose buffer (320 mM sucrose, 5 mM Hepes pH 7.4) supplemented with protease (Roche #05056489001) and phosphatase inhibitors (Invitrogen #A32957) using the Precellys homogenizer (5800 rpm, 2×15 sec with a 30 sec break; soft tissue homogenizing kit CK14 – 2 mL, #P000912-LYSK0-A, Bertin Technologies). Each tissue sample (∼120 mg cortex) was homogenized in 1.5 ml sucrose buffer. Then the homogenates were pooled. Four hippocampi were pooled (∼81 mg) and homogenized in 1.01 ml sucrose buffer. Then, protein homogenates were centrifuged at 1,200 x g for 20 min at 4°C. The resulting pellets were discarded and the supernatants collected and centrifuged at 10,000 x g for 30 min at 4°C. The resulting pellets (P10,000 x g) were then resuspended in sucrose buffer and 1 ml of each membrane pellet was adjusted to 1.8 M sucrose buffer in 5 mM Hepes (pH 7.4) supplemented with protease and phosphatase inhibitors (giving a final volume of 3 ml) and transferred into Beckman coulter ultracentrifuge tubes (#355603). The sucrose gradient was then layered on top of the membrane suspension by 1.33 ml of 1.4 M sucrose buffer with 5 mM Hepes (pH 7.4), 1.33 ml of 1 M sucrose buffer with 5 mM Hepes (pH7.4), 1.33 ml of 0.6 M sucrose buffer with Hepes (pH 7.4) and 1 ml of 320 mM sucrose buffer with Hepes (pH 7.4). The gradient was centrifuged at 285,000g at 4°C for 16 h in a Beckman Optima XPN-90 centrifuge using the 70.1Ti rotor. After centrifugation the gradient was collected in 0.66 ml fractions from top (fraction 1) to the bottom of the tube (fraction 12). The pellet in the tube bottom was resuspended in the last fraction (fraction 12). Equal volumes of each fraction (9.8 µl) were analysed by immunoblotting.

### Ingenuity Pathway Analysis (IPA)

IPA was performed for several sets of differentially altered proteins. In figure 3g, the 9 identified genes shown in figure 3f that are significantly changed both at the transcript and the protein level in 5xFAD brains were examined. In figure 4e, IPA was done for proteins which are concomitantly altered in their abundance both in mouse and human brain tissues. The results of IPA for the differentially expressed proteins across all time points (age 2, 5 and 8) in cortex and hippocampus defined by a “pairwise model” (DE.cortex.Age2, DE.cortex.Age5 and DE.cortex. Age8; DE.hippo.Age2, DE.hippo.Age5 and DE.hippo.Age8) are shown in Additional File 2: Fig. S4a. The results for the DEPs identified through a “full model” (summarized data sets A2TC, A5TC, A8TC and A2TH, A5TH, A8TH) are presented in Additional File 2: Fig. S6a. Finally, in Additional File 2: Fig. S7, it was examined whether IPA pathways are significantly enriched among Aβ-correlated and anticorrelated DEPs.

IPA is based on a library of canonical pathways [49]. The significance of the association between the examined data set and the canonical pathways was measured with a right-tailed Fisher’s exact test to calculate p-values determining the probability that the association between the proteins in the data set and the canonical pathway is explained by chance alone. The p-values were adjusted by the Benjamini-Hochberg multiple testing correction. Only the most significant pathways obtained with each subset of DEPs were depicted (**Fig. 3g, 4e, Additional File 2: Fig. S4a, S6a and S7**).

### Analysis of WT expression profiles

To address the question whether highly expressed mouse brain proteins are more likely to be changed in 5xFAD brains than lowly expressed proteins, we investigated the relationship between WT expression and transgenic effect of the significantly altered proteins (**Additional File 3: Supplementary Excel File 1**) at month 8 in cortex (DE.Cortex.Age8) and hippocampus (DE.hippo.Age8). First, median splits for the wild-type expression values of the proteins identified at month 8 in cortex and hippocampus were carried out in MS-Excel. Groups of low, medium and high WT expression were formed, while the proteins in the medium group were omitted for further analysis. We recognized a correlation, i.e. that highly expressed proteins are more likely to be differentially affected than lowly expressed ones by Aβ aggregates in 5xFAD mouse brains (**Additional File 2: Fig. S3**).

### Gene Ontology enrichment analysis

Six time-resolved data sets of DEPs (months 2, 5 and 8, **Additional File 3: Supplementary Excel File 1**) were selected for a Gene Ontology (GO, RRID:SCR_002811, http://www.geneontology.org/) enrichment analysis (**Additional File 2: Fig. S4b**) that was performed with the Gene Ontology enRIchment anaLysis and visuaLizAtion tool Gorilla [47]. The shared GO term Molecular Function in the DEP data sets was measured. The enrichment of Molecular Function terms in the data sets was computed as an exact p-value of a given minimum hypergeometric (mHG) score, corrected for multiple testing determining the probability that the association between the proteins in the data sets and the total number of proteins that map to the GO term is explained by chance alone (**Additional File 2: Fig. S4b**). The enrichment (**Additional File 2: Fig. S4b**) is defined as (b/n)/ (B/N), where N is the total number of genes; B is the total number of genes associated with a specific GO term; n is the number of genes in the ’target set’ and b is the number of genes in the ’target set’ that are associated with a specific GO term [47].

### Quantification and statistical analysis

Quantification of immunoblots was performed using the iBright Analysis Software (Thermo Fisher Scientific). The scientific data analysis software GraphPad Prism (GraphPad Software, https://www.graphpad.com, RRID:SCR_002798), the R programming environment (R Project for Statistical Computing, RRID:SCR_001905, R Core Development Team, 2006) and MS-Excel were used for data and statistical analysis and graphical representation. Functional analyses of the DEPs were performed using Ingenuity Pathway Analysis [46] and the Gene Ontology enRIchment anaLysis and visuaLizAtion tool Gorilla [47]. Statistical parameters including the value of n (number of independent experimental replications), the definition of precision measures (arithmetic mean ± SD) along with the significance are reported in the figures and figure legends. Data were judged to be statistically significant when p < 0.05 (*), p < 0.01 (**), p < 0.001 (***) and p < 0.0001 (****). Specified statistical tests are described in the method details.

## Results

### Aggregation-prone Aβ42 peptides accumulate faster in hippocampus than in cortex of 5xFAD transgenic mice

Previous investigations have demonstrated progressive accumulation of Aβ42 peptides in brain of 5xFAD mice [21]. Whether accumulation of Aβ42 peptides develops differently in distinct brain regions has not been systematically assessed. Here, we first quantified the abundance of Aβ42 peptides in hippocampus and cortex of 2-, 5- and 8-month-old AD mice and controls using a sandwich ELISA. Brain extracts were supplemented with guanidine HCl to facilitate effective Aβ42 peptide detection in protein samples [48]. We observed a time-dependent increase of Aβ42 peptides in both hippocampus and cortex of 5xFAD tg mice but not in age-matched control animals (**Fig. 1a**). However, Aβ42 peptide levels were significantly higher in hippocampus than in cortex, indicating that the rate of Aβ42 peptide accumulation is distinct in these brain regions. A similar result was also obtained when the abundance of Aβ40 peptides was quantified by ELISA (**Additional File 2: Fig. S1a**). However, it is important to note that in comparison to the measured Aβ42 peptide levels Aβ40 peptides levels are much lower in 5xFAD brains, confirming previously reported results [21].

To assess whether APP and PS1 expression are distinct in hippocampus and cortex of 5xFAD tg mice, we next quantified transcript levels by qPCR. We measured ∼1.5-2-fold higher *APP* and *PSEN1* transcript levels in hippocampus than in cortex (**Additional File 2: Fig. S1b,e**), suggesting that also protein synthesis is higher in hippocampus. This was also confirmed by SDS-PAGE and immunoblotting (**Additional File 2: Fig. S1c,d,f**). Interestingly, the measured Aβ42 peptide levels were ∼3-fold higher in hippocampus than in cortex of 8-month-old tg animals (**Fig. 1a**), suggesting that a ∼1.5-2-fold higher APP and PS1 protein expression in hippocampus compared to cortex leads the massive accumulation of Aβ42 peptides in this brain region. Interestingly, a much higher amount of proteolytically released ∼3 kDa Aβ peptides is present in hippocampus of 8-month-old animals compared to cortex (**Additional File 2: Fig. S1c**). Together, these results suggest that the process of APP cleavage and Aβ42 production is more efficient in hippocampus than in cortex.

### Insoluble Aβ42 aggregates accumulate in larger amounts in hippocampus than in cortex

To quantify the time-dependent formation of insoluble Aβ aggregates in brain, we applied a membrane filter assay (MFA), which facilitates the detection of high-molecular-weight amyloidogenic protein aggregates retained on a cellulose acetate membrane by immunostaining [49]. We found that Aβ aggregate load in 5xFAD mice was significantly higher in hippocampus than in cortex (**Fig. 1b, Additional File 2: Fig. S2a**), suggesting that a large fraction of the Aβ42 peptides detected by ELISA (**Fig. 1a**) are aggregated in AD brains. This is supported by statistical analysis indicating that the time-dependent increase in Aβ42 peptides in 5xFAD brains and the accumulation of Aβ aggregates measured by MFA in hippocampus are positively correlated (**Fig. 1c**). No significant correlation between Aβ42 peptide levels and Aβ aggregate load, however, was obtained for cortical tissue samples (**Additional File 2: Fig. S2b**). Interestingly, in the hippocampus of 5xFAD tg mice, Aβ aggregate load increased ∼10-fold between 5 and 8 months, while no significant increase was observed in cortex (**Fig. 1b**).

Next, we performed a systematic immunohistochemical analysis to investigate the abundance of Aβ amyloid plaques in hippocampus and cortex of 5xFAD tg mice. We applied the Aβ-reactive monoclonal antibody 6E10 (**Additional File 1: Table S1**) and first quantified plaque load in AD and control brains. We detected a significantly higher load (measured plaque volume per tissue volume) in hippocampus than in cortex of 5xFAD tg mice (**Fig. 1d**), essentially confirming the results obtained by MFAs (**Fig. 1b**). No amyloid plaques were detected with the 6E10 antibody in brains of wild-type control mice (data not shown). A similar result was obtained when the number of amyloid plaques was counted in hippocampus and cortex of 5xFAD tg mice (**Fig. 1e**). We also quantified the abundance of Thioflavin S (ThioS)-positive amyloid plaques in AD and control mouse brains. ThioS stains the dense core regions in amyloid plaques, which predominantly contain highly stable fibrillar Aβ42 structures with a typical cross-β structure [50]. Loosely packed “diffuse” plaques detected by immunohistochemical methods are not stained by ThioS (**Fig. 1f**). A comparative analysis revealed slightly fewer ThioS-positive Aβ plaques than 6E10-reactive ones in 5xFAD brains (**Fig. 1e**). In comparison to cortex, many more ThioS-positive Aβ plaques were present in the hippocampus (**Fig. 1e**). Overall, our histological studies revealed that the abundance of Aβ plaques in hippocampus of 5xFAD tg animals was ∼3-5-fold higher than in cortex, indicating that the aggregation rate of Aβ42 peptides is different across brain regions.

Finally, we investigated the intracellular accumulation of Aβ aggregates in 5xFAD brains utilizing an established subcellular fractionation approach [45]. Intracellular membrane fractions were prepared from hippocampal and cortical brain extracts of 8-month-old 5xFAD tg mice by sucrose gradient centrifugation and analysed by SDS-PAGE and immunoblotting (**Additional File 2: Fig. S2c)**. Established marker proteins such as Calnexin (endoplasmic reticulum), Flotillin (lipid rafts) or Lamp1 (lysosomes) were utilized to assess the association of APP and Aβ aggregates with different membrane systems. We observed a co-fractionation of high molecular weight, SDS-stable Aβ aggregates (retained in the stacking gel) with vesicles derived from the ER/endosomal/lysosomal system, indicating that besides extracellular amyloid plaques (**Fig. 1e,f**) also intracellular Aβ aggregates accumulate in 5xFAD brains. As expected, the abundance of Aβ aggregates in hippocampus was higher than in cortex (**Fig. 1g, Additional File 2: Fig S2d,e**). Interestingly, we also observed a co-fractionation of Aβ aggregates with mitochondrial marker proteins (VDAC and NDUFB3), suggesting that also mitochondrial membranes are enriched with our centrifugation procedure. Overall, these biochemical investigations support previously reported immunohistochemical studies, indicating that Aβ42 peptides accumulate in cells of 5xFAD brains [21].

### Overproduction of mutant variants of APP and PS1 leads to time-dependent proteome changes in hippocampus and cortex of 5xFAD tg mice

To assess whether APP and PS1 overproduction in brains of 5xFAD tg mice is accompanied by global protein expression changes, we performed a quantitative mass spectrometry (MS)-based proteomics analysis. Protein abundance was measured by label-free quantification (LFQ) in hippocampal and cortical tissues prepared from 2-, 5- and 8-month-old 5xFAD and control mice. By using the quantitative readouts obtained from MaxLFQ analysis [51], we identified ∼5,000 proteins per tissue on average. We utilized a rather conservative approach to analyse the proteomics data. No imputation was done on missing LFQ intensity values.

First, we compared AD genotypes at indicated ages with their wild-type counterparts (pairwise model). We identified large numbers of differently expressed proteins (DEPs) in both hippocampal and cortical tissues (**Fig. 2a,b, Additional File 3: Supplementary Excel File 1**). The global proteome response was strongest at 8 months, when deposition of Aβ aggregates was readily detectable in mouse brains (**Fig. 1d**), indicating that aggregate formation is associated with perturbation of the mouse brain proteome. In hippocampus, e.g., 57 proteins were differentially expressed at 5 months, while 544 proteins were changed at 8 months of age (**Fig. 2b**). Overall, we identified similar numbers of up- and down-regulated proteins in hippocampal and cortical tissues (**Fig. 2b**), although in the cortex at 2 months of age the number of upregulated proteins was ∼7-fold higher than the number of downregulated ones (**Fig. 2b, Additional File 3: Supplementary Excel File 1**).

To assess whether Aβ aggregation predominantly influences the abundance of highly expressed proteins, we utilized the protein abundance measurements in brains of wild-type mice to define groups of proteins with high and low protein expression. Next, we determined whether protein abundance changes detected in 5xFAD brains are more frequent among the highly expressed mouse proteins than among the lowly expressed ones. Interestingly, we found a significantly higher number of Aβ-associated protein changes among the highly expressed mouse brain proteins (**Additional File 2: Fig. S3**), supporting our hypothesis that Aβ aggregation perturbs higher abundant proteins with a higher likelihood than lower abundant proteins.

We performed IPA (ingenuity pathway analysis) for an unbiased assessment of whether observed proteome changes reflect alterations in specific cellular processes and/or pathways. This analysis revealed that many proteins involved in oxidative phosphorylation and mitochondrial functions change in their abundance over time in AD mouse brains (**Additional File 2: Fig. S4a**), while changes in proteins involved in phagosome maturation and synaptogenesis were predominantly detected relatively late in the disease process (8 months), when large amounts of Aβ42 aggregates are already present in AD brains. This was also confirmed by gene ontology (GO) term enrichment analysis (**Additional File 2: Fig. S4b**).

Finally, we assessed whether the observed protein abundance changes in AD brains originate from different cell types. To address this question the enrichment of cell type-specific marker proteins, which were previously defined for astrocytes, microglia, neurons and oligodendrocytes, was investigated [44]. We found only a small number of marker protein changes among the abnormally up-and down-regulated proteins in 5xFAD brains (**Additional File 2: Fig. S5**). However, in hippocampal tissues of 8-month-old animals multiple neuronal marker proteins were significantly changed, suggesting that the observed global proteome response at least in part originates from Aβ-perturbed neurons.

### The proteome response in cortical and hippocampal tissues of 5xFAD brains is distinct

In order to compare proteome changes between age groups and different tissues, we have limited the differential analysis to proteins reliably measured in each condition (genotype, age and tissue) to ensure that the transgene effect difference between conditions can be statistically estimated (for details see Methods section). Simple pairwise comparisons would be insufficient because they do not allow the computation of interaction effects. However, this requirement considerably limits the number of proteins that can be modelled: only 2761 (∼55% of the originally identified proteins) fulfil the completeness requirements across all conditions.

Utilizing the 2761 proteins in the “full model”, we identified 131 differentially expressed proteins (DEPs) in cortex (2, 5 and 8 months), while 558 DEPs were detected in hippocampal tissues (**Fig. 2c, Additional File 3: Supplementary Excel File 2**). Thus, in hippocampal tissues where the time-dependent accumulation of Aβ aggregates is more rapid than in cortical tissues (**Fig. 1a**), a ∼4-fold higher number of DEPs was observed, suggesting that the proteome response in these brain regions is different.

Next, we focussed on changes in DEPs between time points and tissues. We calculated significant differential protein abundance changes between ages 5 and 2 (A52), 8 and 5 (A85) and 8 and 2 (A82) months for both cortical (C) and hippocampal (H) tissues (T) (**Fig. 2c**). As expected, in comparison to cortical tissues the number of significant differential protein abundance changes was ∼3-fold higher in hippocampal tissues, confirming our initial observations that increased deposition of Aβ aggregates in brain is associated with higher numbers of DEPs (**Fig. 2c**). The number of significant differential protein abundance changes was higher between 8 and 2 months (A82TH) than between 5 and 2 (A52TH) or 8 and 5 (A85TH) months, supporting our hypothesis that transgene expression leads to a progressive proteome response that is more pronounced after a time period of 6 (A82) than of 3 (A52 or A85) months. Importantly, differential protein abundance effects were also observed when cortical and hippocampal tissues prepared at 2, 5 and 8 months (A2THC, A5THC and A8THC) were directly compared with each other (**Fig. 2c**), substantiating our view that distinct cellular pathways are altered in both tissues upon transgene expression.

In order to identify proteome changes that occur both in cortical and hippocampal tissues, we next looked at overlapping and non-overlapping DEPs in these brain tissues. We found a relatively small fraction of proteins (56 proteins, 8.8%) that were dysregulated upon transgene expression in both hippocampus and cortex (**Fig. 2d**). A similar result was obtained when proteins, which are significantly increased or decreased in their abundance in hippocampal and cortical tissues, were assessed for overlapping and non-overlapping dysregulated proteins (**Fig. 2e, f**).

Thus, our analysis indicates that most protein changes observed in hippocampus and cortex are distinct, suggesting that very different cellular pathways and/or processes are altered in these brain regions. This view is also supported by ingenuity pathway analysis (IPA), demonstrating that mostly different cellular pathways are dysregulated over time in these brain regions (**Additional File 2: Fig. S6a**). We observed dysregulation of mitochondrial processes and mTOR signaling predominantly in cortical tissues, while synaptic processes including long term potentiation and axon guidance signaling were changed in hippocampal tissues (**Additional File 2: Fig. S6a**). Pathways altered both in hippocampus and cortex included oxidative phosphorylation, transcription regulation, DNA methylation and repair, indicating that important cellular functions are changed in both tissues. Time-dependent dysregulation of synaptic, lysosomal and mitochondrial proteins in hippocampal and cortical tissues was confirmed by gene ontology (GO) term enrichment analysis (**Additional File 2: Fig. S6b**).

To investigate the relationships between protein changes observed in hippocampal and cortical tissues at 2, 5 and 8 months (**Additional File 3: Supplementary Excel File 2**), we calculated Spearman correlation coefficients, which can be used to assess the similarity between protein expression data sets. We observed strong correlations, when expression changes of 5- and 8-month-old (A5TH vs. A8TH) hippocampal tissues were compared (**Fig. 2g**). Similarly, a strong positive correlation was obtained, when expression changes in 8-month-old cortical tissues (A8TC) were compared with expression changes in 5- (A5TH) and 8-month-old (A8TH) hippocampal tissues. This indicates that cortical and hippocampal tissues that contain Aβ42 aggregate deposits (**Fig. 1d-f**) exhibit at least in part a similar protein response. Interestingly, strong anticorrelations were obtained when protein expression changes of 2- and 8-month-old animals were compared (**Fig. 2g**), indicating that the proteome response in young AD tg mice is the opposite compared to older mice that progressively accumulate Aβ42 aggregates. The anticorrelation of protein changes in the cortex of 2- and 8-month-old animals is exemplarily shown in **Fig.2h**. Interestingly, in cortical tissues of 2-month-old animals, we observed a strong increase in the abundance of mitochondrial proteins that play a key role in oxidative phosphorylation and ATP production (**Fig. 2i**), suggesting that expression of mutant human *APP* and *PSEN1* in neurons of young mice is associated with increased mitochondrial activity and ATP production. However, this proteome response gets lost over time when the mice get older concomitantly with the accumulation of Aβ42 aggregates, supporting the hypothesis that progressive Aβ aggregation in neurons impair critical mitochondrial functions [52]. Finally, we quantified mRNA changes in cortical and hippocampal tissues and assessed whether the observed time-dependent changes of mitochondrial proteins (**Fig. 2i**) is also observed at the transcript level. Except for one gene, we found no significant transcript changes, indicating that the observed mitochondrial protein abundance changes (**Fig. 2i**) most likely are not caused by transcriptional dysregulation.

Overall, these investigations indicate that the proteome response in hippocampus and cortex of *APP* and *PSEN1* expressing mice is largely distinct. However, in older brains (5 and 8 months), when insoluble Aβ42 aggregates are readily detectable, proteome changes in hippocampal and cortical tissues appear to be more similar (**Fig. 2g**), suggesting that intracellular and/or extracellular Aβ “aggregate stress” perturbs similar pathways/processes in different brain regions.

### Identification of Aβ-correlated and anticorrelated protein alterations in 5xFAD brains

To explore molecular events associated with progressive Aβ accumulation, we identified protein abundance changes that are correlated or anticorrelated to the time-dependent accumulation of Aβ aggregates in hippocampus and cortex of 5xFAD transgenic mice. In total, we identified 699 DEPs (pairwise model, **Fig. 2a, b**), of which 182 (26.0%) were correlated and 148 (21.2%) were anticorrelated (**Additional File 3: Supplementary Excel File 3**) to the aggregate load in both brain regions (**Fig. 3a, b**). As expected, in hippocampal tissue, which contains significantly higher levels of Aβ aggregates than the cortex (**Fig. 1d-g**), a significantly higher number of correlated and anticorrelated protein changes was found.

Interestingly, we observed a strong positive correlation between the most influential AD risk factor apolipoprotein (ApoE) and the time-dependent accumulation of Aβ aggregates in both hippocampal and cortical tissues (**Fig. 3b**), suggesting that this protein gets upregulated in response to Aβ aggregate deposition. Our pathway analysis (IPA) revealed a significant activation of the LXR/RXR pathway in hippocampal tissues of 5xFAD tg animals (**Additional File 2: Fig. S4a**). Liver X receptors (LXRs) and retinoid X receptors (RXRs) were reported to function as transcription factors regulating cholesterol and fatty acid homeostasis [53] including the transcriptional activation of the apolipoprotein E (*APOE*) gene [54]. This suggests that the observed activation of the LXR/RXR pathway in 5xFAD brains is accompanied by an upregulation of *APOE* gene expression. To address this question, we quantified mRNAs by RNAseq and found that *APOE* transcript levels in hippocampal and cortical tissues (5 months) of 5xFAD tg mice are indeed increased compared to wild-type controls (**Fig. 3c**). However, in 8-months-old animals *APOE* transcript levels were decreased in cortical but not in hippocampal tissues, suggesting that at a later stage transcriptional regulation of *APOE* expression is distinct in these tissues. Strikingly, a very pronounced time-dependent increase of ApoE protein levels was detected in both brain regions at 5 and 8 months, indicating that *APOE* transcript and protein levels in cortical tissues of 8 months-old animals are not well correlated (**Fig. 3d**).

Also, we found that the proteins C1qb, Hexb and Acat2 are strongly correlated with the appearance of Aβ aggregates in both brain regions in 5xFAD mice (**Fig. 3b**). Strikingly, all these proteins have been previously linked to AD as well as to cellular processes that control the progressive accumulation of Aβ aggregates [17, 55]. C1qb is the initiating protein of the classical complement cascade, which associates with Aβ pathology in patient brains and was previously shown to mediate synapse loss in transgenic AD models [56]. The gene encoding Hexb was previously identified as a putative AD risk gene. It is important for lysosome function and influences Aβ accumulation in mouse brains [57]. Acat2 is central to lipid metabolism and inhibitors of this protein were shown to modulate Aβ production and reduce Aβ accumulation in a transgenic model of AD [58]. In contrast to these positively correlated proteins, we found Arpc5 and Glrx to be significantly anticorrelated to Aβ-accumulation in both brain regions (**Fig. 3b**), suggesting that their activity gets decreased upon progressive Aβ aggregation in 5xFAD brains. Arpc5 is a subunit of the Arp2/3 complex that promotes actin filament assembly [59] and is known to influence neurite outgrowth. A direct link to AD has not been described, however, it is well-known that progressive Aβ accumulation perturbs actin dynamics in neuronal cells [60]. Glrx, a member of the glutaredoxin family, was previously shown to influence cognitive deficits in an AD mouse model [61].

Finally, we performed IPA to elucidate sub-cellular pathways that are potentially influenced by progressive Aβ42 accumulation in brains of 5xFAD transgenic mice. We observed that multiple proteins involved in synaptogenesis signaling and phagosome maturation are altered significantly in their abundance in hippocampal tissues of 5xFAD (**Additional File 2: Fig. S7**), suggesting that progressive Aβ42 accumulation in this brain region perturbs critical synapse functions. This is in agreement with previous observations indicating that small Aβ42 assemblies, often termed oligomers, target synapses and can perturb the activity of specific synaptic proteins [18, 62]. Interestingly, in cortical tissues, where the time-dependent deposition of Aβ42 aggregates is significantly lower compared to hippocampal tissues (**Fig. 1a-e**), we observed no synaptic protein changes (**Additional File 2: Fig. S7**), supporting our hypothesis that Aβ42 aggregation and perturbation of protein homeostasis at synapses are linked. Overall, these investigations indicate that progressive accumulation of Aβ42 peptides in AD mouse brains is associated with alterations in synaptic functions, which may lead to neuronal dysfunction and memory impairment.

### Aβ-correlated and anticorrelated proteome and transcriptome changes are largely distinct in 5xFAD brains

We generated mRNA expression data sets by RNAseq (**Additional File 3: Supplementary Excel File 4)** to compare proteome and transcriptome changes in hippocampal and cortical tissues of 2-, 5- and 8-month-old 5xFAD transgenic mice and age-matched wild-type controls. Here, we specifically focused on transcripts that encode proteins whose abundance is significantly changed in 5xFAD brains and that are also correlated or anticorrelated to the Aβ aggregate load. As expected, we detected transcripts for all of the 329 significantly dysregulated Aβ-correlated and anticorrelated proteins in 5xFAD brains (**Fig. 3e, Additional File 3: Supplementary Excel File 4**). However, only a small fraction of these transcripts (2.8%) was significantly changed in 5xFAD brains (**Fig. 3e**), indicating that most of the protein abundance changes associated with Aβ aggregation in 5xFAD brains are caused by posttranscriptional mechanisms rather than transcriptional dysregulation. We identified 9 genes that are significantly changed both at the transcript and the protein level in 5xFAD brains. (**Fig. 3f**). Interestingly, a large fraction of these genes encodes proteins that play a functional role in the complement system (C1qa, C1qb and C1qc), suggesting that transcriptome changes predominantly drive microglia activation and inflammatory processes in 5xFAD brains. This is also supported by IPA, indicating that proteins involved in the complement system are overrepresented among the transcriptionally dysregulated proteins (**Fig. 3g, Additional File 3: Supplementary Excel File 5**). However, it is important to note that also the abundance of proteins with specific functions in lysosomes (Hexb), metabolic processes (Acat2) and protein degradation pathways (Ctsd, Cst3) are transcriptionally regulated in 5xFAD brains.

### A fraction of dysregulated proteins in 5xFAD brains is concordantly altered in postmortem brains of AD patients

Next, we assessed whether proteins that change in their abundance in brains of 5xFAD mice (**Fig. 2**) overlap with proteome changes detected in postmortem brains of AD patients. For our comparisons, we focussed on two recently reported large-scale proteomics studies (J20 and J22) that measured the abundances of human proteins in postmortem brains of AD patients and controls using different quantitative mass spectrometry approaches [63, 64]. In total, in J20 and J22 3950 significantly altered AD-associated human proteins were identified, of which 689 proteins (17.4%) were identified in both studies. In addition, we compared our DEPs from mouse brains with a recently reported amyloid plaque proteome (termed D22; 816 proteins), which was obtained by label free MS analysis of microdissected Aβ plaques [65]. Here, we focused our efforts on protein abundance changes detected in 8-month-old hippocampal and cortical mouse tissues, which contain high levels of potentially proteotoxic Aβ42 aggregates (**Fig. 1d, e**). In addition, protein abundance changes in 5-month-old hippocampal tissues, which also contain significant amounts of amyloidogenic protein aggregates, were compared with patient protein measurements. We found a substantial overlap between mouse **(Additional File 3: Supplementary Excel File 1)** and human DEPs (**Fig. 4a**), indicating that 5xFAD tg mice at least in part recapitulate the proteomic changes observed in AD patient brains. Interestingly, a small number of proteins such as ApoE, Clu or NDUFA12 were significantly altered in all investigated data sets, suggesting that they are potential marker proteins that robustly reflect the amyloid state both in patient and mouse brains.

To assess whether the observed protein changes in mouse and patient brains are similar, we calculated Spearman correlation coefficients for all pairwise data cross comparisons (**Fig. 4b, c**). Strikingly, we observed a strong correlation between the protein measurements in hippocampal or cortical tissues of 5xFAD mice and the protein measurements in patient amyloid plaques (D22 data set), indicating that in 5xFAD brains multiple potentially disease-relevant protein changes are recapitulated that are also detectable in amyloid plaques of patient brains [65]. Also, moderate positive correlations were obtained when protein measurements of cortical (8 month) or hippocampal (5 month) tissues were compared with the large-scale proteomics data sets (J20 or J22), supporting our hypothesis that disease-relevant amyloid-related proteome changes are recapitulated in 5xFAD mice.

Interestingly, our analysis suggests that protein changes that are measured in mouse brain tissues with a relatively low amount of Aβ aggregates (e.g., 5-month-old hippocampal tissues or 8-month-old cortical tissues) are better predictors of the disease state in AD patients than protein changes measured in 8-month-old hippocampal tissues, which contain a high amount of insoluble Aβ aggregates (**Fig. 1d**). As expected, strong positive correlations were obtained when protein measurements of different human brain samples were compared in a pairwise manner (e.g., D22 with J22; **Fig. 4b**).

Next, utilizing the results from all data cross comparisons, we generated a data set that exclusively contains proteins, which are concomitantly altered in their abundance both in mouse and human brain tissues. In total, 140 potential AD-relevant protein changes were defined (**Additional File 3: Supplementary Excel File 6**), of which 67 (47.9%) were abnormally up- and 73 (52.1%) were downregulated in hippocampal and/or cortical mouse tissues of 5xFAD brains. The top 15 proteins with the highest fold changes from this data set were selected in order to define potentially patient-relevant mouse protein signatures (Sig+ and Sig-; **Additional File 2: Fig. S8, Additional File 3: Supplementary Excel File 6**). As shown in **Fig. 4d**, proteins such as ApoE, Clu and GFAP are strongly up-regulated both in 5xFAD and AD patient brains, while proteins such as CNP, NDUFA12 and MBP are downregulated. Finally, we performed IPA using the data set of concordantly changed mouse proteins (**Additional File 3: Supplementary Excel File 6)**. Strikingly, proteins that play a key role in mitochondria and synapses were enriched in this data set (**Fig. 4e, Additional File 3: Supplementary Excel File 5**), supporting our hypothesis that “Aβ aggregate stress” perturbs important mitochondrial and synaptic functions in mouse and human AD brains.

### The levels of the Arf-like GTPase Arl8b are abnormally increased in hippocampal tissues of 5xFAD mice

In its earliest clinical phase, AD is characterized by memory impairment [66], suggesting hippocampal synaptic failure is a critical early event in pathogenesis.

Therefore, we next searched for neuronal proteins in the proteomics data set that are strongly altered in their abundance, (2) expressed in hippocampal tissues and (3) known to influence synaptic processes in neurons. Interestingly, we found that, among the Aβ42 aggregate-correlated proteins with the highest p-value, the ARF-like GTPase Arl8b is significantly increased in hippocampal tissues (**Fig. 3b,5a**). Also, this protein was found to be concordantly altered both in 5xFAD and postmortem brains of AD patients (**Fig. 4a-d**), suggesting that its protein changes in mouse brains might have predictive value for the pathological situation in AD patient brains. Arl8b was reported to associate with lysosomes and motor proteins in axons [67, 68] and to control the anterograde transport of synaptic vesicles, which is critical for efficient synapse function [69]. Thus, we hypothesized that a specific investigation of Arl8b in the context of AD might provide novel insights into how the formation of Aβ42 assemblies in hippocampal tissues leads to lysosomal dysfunction and/or synaptic failure and memory impairment.

To independently validate the changes in Arl8b abundance initially observed with mass spectrometry (**Fig. 5a**), we analyzed protein extracts prepared from hippocampal tissues of 8-month-old 5xFAD and control mice by SDS-PAGE and immunoblotting. We confirmed that protein levels of Arl8b in AD mouse brains are significantly increased compared to age-matched controls (**Fig. 5b,c**).

Next, we assessed the abundance of Arl8b in 5xFAD brains by immunohistochemistry. Strikingly, we observed a time-dependent accumulation of Arl8b in ring-like structures surrounding Aβ42 amyloid plaques (**Fig. 5d** **and Additional File 2: Fig. S9**). These structures have been reported to be axonal lysosome accumulations formed in close vicinity to amyloid plaques in AD brains [70]. To determine whether the observed Arl8b-postive structures are indeed lysosomal accumulations, we stained brain slices with an antibody that recognizes the lysosomal marker protein Lamp1 ([71], **Additional File 1: Table S1**). As expected, we observed co-staining of Arl8b and Lamp1 in 5xFAD brains (**Fig. 5e**). The formation of Aβ aggregates in mouse brains was confirmed with the monoclonal antibody 352 (**Fig. 5e, Additional File 1: Table S1**), which specifically recognizes Aβ42 fibrillar aggregates but not soluble monomers *in vitro* [30].

Finally, we assessed the relationship between the volume of Aβ42 plaques in 2-, 5- and 8-month-old 5xFAD mice and the volume of Arl8b-positive lysosomal structures accumulating in their vicinity. We found a strong correlation between Arl8b- and 6E10-stained structures in hippocampal tissues of all three ages (**Fig. 5f**), suggesting that the time-dependent formation of amyloid plaques determines the size of the adjacent Arl8b-postive lysosomal structures in 5xFAD brains. An association of Arl8b with lysosomal membranes and proteins has previously been described [67, 68], supporting our immunobiological results.

## Arl8b protein levels are abnormally increased in postmortem brain tissues and CSF samples of AD patients

To analyze whether Arl8b accumulates in AD patient brains, we investigated protein extracts prepared from postmortem brain tissues of AD patients and age-matched controls (10 each) using a membrane filter assay (MFA) [49]. With this method, large protein aggregates, e.g., Aβ and tau assemblies, retained on a cellulose acetate membrane after filtration (pore size 0.2 μm), can be detected by immunoblotting [30, 36]. We found that the immunoreactivity of Arl8a and Arl8b, both specifically recognized by the anti-Arl8a/b antibody (**Additional File 1: Table S1**), is significantly increased in AD patient brains compared to age-matched controls (**Fig. 6a,b**), indicating that the abundance of these functionally and structurally closely related proteins is abnormally elevated in AD patient brains. A similar result was obtained with an Arl8b-specific antibody (**Additional File 2: Fig. S10a, b**).

**Figure 6.**
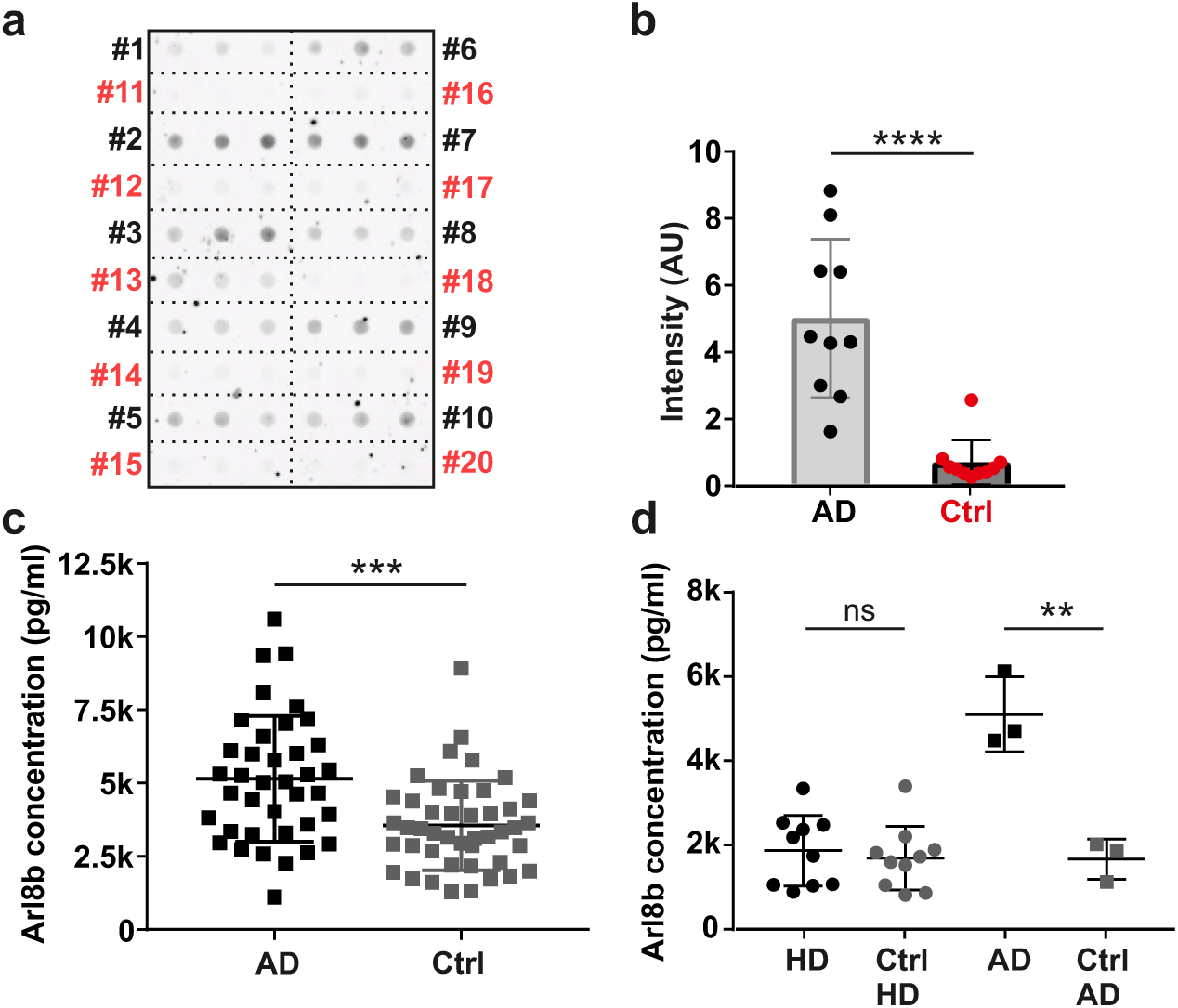
Analysis of Arl8b expression in human brain and CSF samples. **(a)** Detection of Arl8 protein aggregates in postmortem brain homogenates of 10 AD patients (1 to 10, black lettering) and 10 age-matched controls (11 to 20, red lettering) using a native MFA. Triplicates per sample were filtered. For immunoblotting an anti-Arl8a/b antibody was used. **(b)** Quantification of protein aggregates retained on filter membrane in (**a**) was performed using Aida image analysis software. Data are expressed as mean ± SD. The statistical significance was assessed with an unpaired, two-tailed t test (****, p < 0.0001). **(c)** Determination of Arl8b concentrations in CSF samples of 38 AD patients and 44 control individuals (Ctrl) using an ELISA. Data represent mean ± SD. Statistical significance was determined using an unpaired, two-tailed t test (***, p = 0.0002). **(d)** Arl8b ELISA using CSF samples of 10 Huntington’s Disease (HD) patients, 10 controls (Ctrl HD), 3 AD patients and 3 controls (Ctrl AD). Data are mean ± SD. Statistical significance was evaluated using an unpaired, two-tailed t test between HD and Ctrl HD, and AD and Ctrl AD groups (**, p = 0.0041).

Finally, we investigated the levels of Arl8b in cerebrospinal fluid (CSF) of AD patients and control individuals using an ELISA. The demographic, clinical and CSF biomarker characteristics of AD patients and control individuals are summarized in **Additional File 1: Table S2**. Strikingly, we measured significantly higher Arl8b protein levels in CSF samples of AD patients compared to controls (**Fig. 6c**), confirming that protein abundance is abnormally increased in AD brains. In strong contrast, no significant increase of Arl8b protein levels was measured when CSF samples obtained from HD patients and controls were analyzed with ELISA (**Fig. 6d**), indicating that the observed increase in protein abundance is an AD-specific phenomenon. We next assessed the correlations of Arl8b protein levels with other well-established AD biomarkers such as Aβ(1-42), Aβ(1-40), Aβ42/Aβ40, t-tau and p-tau [72]. Spearman correlation analysis revealed a significant moderate negative correlation between Arl8b protein levels and CSF Aβ42/Aβ40 ratios (*r_S_*=-0.405; p=0.0002), supporting our hypothesis that increased Arl8b protein levels are associated with the deposition of Aβ42 protein aggregates in patient brains (**Additional File 1: Table S3**). A weak negative correlation was also observed with Aβ(1-42) CSF peptide levels, while no significant correlation was obtained with Aβ(1-40) levels. Interestingly, we also observed a moderate positive correlation between Arl8b and p-tau protein levels in CSF samples, suggesting that an increase of Arl8b is also associated with cognitive impairment in AD patients. This is also supported by total tau (t-tau) measurements, which show a weak correlation to Arl8b protein levels. Overall, these studies suggest that Arl8b might have potential as a novel CSF biomarker for monitoring the abnormal accumulation of lysosome structures in AD patient brains.

## Discussion

The progressive deposition of Aβ42 peptides in amyloid plaques and intraneuronal structures in patient brains is a pathological hallmark of AD [73]. Aβ aggregates in patient brains are detected before the onset of neurodegeneration and memory impairment in AD patients [74], indicating that their formation is an early event in pathogenesis. However, it was shown reproducibly that the formation of Aβ plaques in patient brains is not a good predictor of neurodegeneration [75]. This suggests that Aβ aggregation is relevant but not solely responsible for disease onset and that other molecular changes drive progressive neurodegeneration in patient brain. Evidence was reported that Aβ aggregation in disease models triggers tau accumulation, which is a good predictor of neurodegeneration and memory impairment [76, 77]. Thus, accumulation of Aβ peptides in patient brains is broadly regarded as a key early event in pathogenesis, leading to downstream neuropathological changes such as the accumulation of tau or abnormal microglia activation and eventually to neurodegeneration.

Here, we report protein abundance changes associated with the progressive accumulation of Aβ42 aggregates in brains of 5xFAD tg mice (**Fig. 2a-i** and **Fig. 3a-f**). 5xFAD tg mice are a well-characterized disease model expressing two human genes (*APP* and *PSEN1*) with familial AD mutations in neurons, leading to the rapid accumulation of intra- and extracellular Aβ42 aggregates in brains [21]. Progressive deposition of Aβ42 structures is accompanied by microglia activation, a change of synaptic proteins and neurodegeneration, suggesting that Aβ42 aggregation in 5xFAD mouse brains drives important downstream proteomic changes that are also observed in AD patient brains. Various proteomics and transcriptomics studies have uncovered that microglial phagocytosis is significantly perturbed in 5xFAD brains, suggesting that extracellular Aβ42 aggregation predominantly leads to an inflammatory response that over time is associated with neuronal dysfunction and toxicity [78]. However, it is unclear whether proteome changes that accompany the accumulation of intra- and extracellular Aβ42 aggregates in 5xFAD are conserved in patient brains.

Based on our comparative analyses, (**Fig. 4a-d**), we suggest that progressive Aβ aggregation in 5xFAD and AD patient brains may lead to similar perturbations of the proteome and could provide valuable information about critical pathways perturbed in disease. We hypothesize that the proteome response to Aβ-aggregate stress is conserved in mouse and human brains to a considerable extent and at least a fraction of the proteome changes observed in patient brains may be caused by progressive Aβ aggregation. Interestingly, our proteome-based data cross comparisons indicate that protein changes in mouse brain tissues with a relatively low amount of Aβ aggregates are more similar to protein changes in AD patient brains than protein changes in tissues, which contain a high Aβ aggregate load (**Fig. 4b**). This suggests that overexpression of mutant variants of *APP* and *PSEN1* in hippocampal neurons of 5xFAD mice after some time (8 months) leads to a proteome response that is distinct from the situation in patient brains. Together these studies indicate that 5xFAD transgenic animals clearly recapitulate important features of the pathogenic process in patient brains. However, timing and brain region-specific investigations are critical for the detection of potential disease-relevant proteome changes in mouse brains. We propose that the proteome changes measured in the hippocampus of 5-month-old 5xFAD mice recapitulate the disease situation in AD patient brains better than the protein changes measured in 8-month-old animals, where massive amounts of Aβ42 aggregates are detectable by immunohistochemistry or biochemical methods. In comparison, the situation is different when proteome changes in cortical tissues of 5xFAD mice are assessed where the rate of Aβ42 aggregate accumulation is significantly slower than in hippocampal tissues (**Fig. 1d**). Thus, our investigations suggest that 5xFAD mice with their massive accumulation of Aβ42 aggregates in a relatively short time (2-8 months) are a model of established rather than asymptomatic AD. We propose that pharmacological interventions in 5xFAD mice may have a predictive value for AD patients, if conserved Aβ42 aggregate-associated proteome changes could be reversed by drug treatment. Previously, small molecules and therapeutic antibodies that directly target Aβ aggregate assemblies and reduce their propagation in model systems have been described [30, 79]. Whether their application, might reverse the proteome changes in 5xFAD brains defined here would need to be assessed.

We provide evidence that progressive Aβ42 aggregation is associated with a reduced abundance of multiple mitochondrial proteins (**Fig. 2i**), suggesting that Aβ42 accumulation drives mitochondrial dysfunction in 5xFAD brains. Strikingly, a loss of mitochondrial proteins is also a characteristic feature of AD brains [80], supporting the hypothesis that decreased energy production is associated with cognitive decline in AD patients [81]. This view is supported by a recent large-scale proteome study, indicating that a stable cognitive trajectory in advanced age is associated with higher levels of mitochondrial proteins and increased neuronal mitochondrial activity [82]. Enhancing mitochondrial proteostasis with pharmaceutical compounds reduced Aβ aggregate-induced proteotoxicity in model systems [19], substantiating the link between Aβ pathology and mitochondrial dysfunction. The underlying molecular mechanism by which the Aβ42 peptides in 5xFAD brains perturb the mitochondrial proteome currently remains unclear, however. We found here that both APP and insoluble Aβ42 aggregates indeed co-fractionate with mitochondrial membranes in 5xFAD brains, suggesting that they also might interact with mitochondria (**Fig. 1g**). It was previously reported that APP and its cleavage products accumulate in mitochondria [83, 84], suggesting they might directly perturb the electron transport machinery, thereby decreasing ATP synthesis [85]. However, toxicity might also be caused by Aβ42 assemblies acting on regulators of gene transcription such as H2AFZ (**Fig. 3b**) or on the surface of mitochondria, perturbing the import of critical mitochondrial proteins or influencing fission, transport or degradation of mitochondria [81]. Further mechanistic investigations are clearly necessary to elucidate the link between Aβ42 aggregation and mitochondrial dysfunction in 5xFAD brains and other model systems. Besides oxidative phosphorylation in mitochondria, various pathways and subcellular processes such as carbohydrate metabolism or synaptogenesis are potentially perturbed in response to Aβ42 accumulation in 5xFAD brains (**Fig. 4e** and **Additional File 2: Fig. S7**). The impact of Aβ42 overproduction on brain physiology is likely highly complex and brain region specific. Links between energy metabolism, synaptic transmission and inflammation have been described in the aging brain and in AD [86, 87], suggesting that the generation of multiscale causal networks and the integration of additional information such as protein-protein interaction data is necessary to elucidate the specific role of Aβ aggregates in mouse brains.

Our studies indicate that progressive Aβ42 aggregation leads to the time-dependent accumulation of the small GTPase Arl8b in brains of 5xFAD mice (**Fig. 5a-f**). This protein mediates axonal transport of both lysosomes and synaptic vesicles in neurons [88, 89], suggesting that its function is of critical importance for the maintenance of synaptic compartments and neurotransmission. This is supported by gene knock-down experiments in hippocampal neurons indicating that loss of Arl8b leads to the accumulation of synaptic vesicles (SVs) in neuronal cell bodies and the depletion of active zone proteins from presynaptic sites [69]. Our studies showed that Arl8b as well as lysosomal proteins such as LAMP1 are enriched in the vicinity of extracellular amyloid plaques, indicating that progressive Aβ42 aggregation is critical for the massive accumulation of lysosomal structures in 5xFAD brains (**Fig. 5d-f**).

This is in agreement with previous investigations demonstrating that protease-deficient axonal lysosomes accumulate in the vicinity of amyloid plaques [70]. Recently, an accumulation of the GTPase Arl8b in the vicinity of amyloid plaques in patient brains has also been demonstrated by immunohistochemistry [65]. Its abnormal accumulation in axons could lead to a perturbation of vesicle transport processes, which often is accompanied with synaptic dysfunction and alteration of synaptic transmission. However, Arl8b accumulation may also stimulate lysosome exocytosis that is known to affect APP processing and Aβ secretion [90]. An increase of lysosomal exocytosis has been described in cancer cells overproducing Arl8b, hinting that this process may also be enhanced in neuronal cells, which contain lysosome-like organelles with high local concentrations of Arl8b (**Fig. 5d**). However, the functional consequences of Arl8b accumulation in axonal lysosomes in the presence of amyloid aggregates need to be studied further in model systems.

Interestingly, our investigations indicate that Arl8b protein levels are significantly increased both in CSF and postmortem brain tissues of AD patients (**Fig. 6a-d**), suggesting that this protein may have the potential to be utilized as a biomarker to monitor lysosomal perturbations in AD. Cell-based experiments have shown that lysosomal cathepsins play an important role in the generation of Aβ peptides [91]. Also, lysosomal membrane alterations have been observed in brains of AD patients [92], and accumulation of Aβ42 in neuronal cells was shown to lead to changes in the abundance of lysosomal organelles and cell death [93]. Thus, it seems adequate to speculate that lysosome-associated proteins that change in their abundance in AD brains, are putative biomarkers that might predict disease development. Currently, the well-established pathological polypeptides such as total-tau, phospho-tau and Aβ42 are measured in CSF to detect incipient and established AD in patients [72, 94]. However, major research efforts are on the way to find additional fluid biomarkers to predict disease onset and monitor progression more accurately [63, 64, 80, 94–96]. Previous studies have shown, e.g., that neurofilament light chain (NfL) and neurogranin are promising AD biomarkers [97, 98]. We envision that a better definition of AD will clearly require a more accurate characterization and understanding of the sequence of events that lead to cognitive impairment. The measurement of Arl8b protein levels in CSF or plasma of AD patients might help to meet this goal. However, more comprehensive studies with symptomatic and asymptomatic AD patients and controls will be necessary to further assess the predictive power of Arl8b as a clinical biomarker.

## Conclusions

Our studies indicate that Aβ42 driven aggregate formation is associated with time-depended and brain region-specific proteome changes in 5xFAD mouse brains. We detected 329 dysregulated proteins correlating or anticorrelating with Aβ aggregation in hippocampus and cortex of 5xFAD mice. Most of these protein changes were caused by posttranscriptional mechanisms, only a minor part was associated with transcriptional dysregulation. A fraction of the Aβ-correlated and anticorrelated DEPs was conserved in postmortem brains of AD patients revealing that proteome changes in 5xFAD mice recapitulate disease-relevant changes in AD patient brains. Among the group of Aβ42-correlating proteins, we have found the lysosome-associated protein Arl8b, which is present in increased levels in CSF samples of AD patients and might have potential as an AD biomarker.

## Additional Files

### Additional File 1

**Table S1.** Commercial antibodies used in this study. **Table S2.** Characteristics of AD and HD patients and corresponding controls. **Table S3.** Spearman correlation analysis of Arl8b protein level measurements and AD biomarker levels.

### Additional File 2

**Fig. S1**. Aβ peptide levels and *APP* and *PSEN1* expression in hippocampus and cortex of 5xFAD mice. **Fig. S2**. Analysis of Aβ aggregate formation using membrane filter assays (MFA) and sucrose gradient centrifugations. **Fig. S3**. Analysis of wild-type expression profiles to assess whether the protein abundance changes detected in 5xFAD brains are more frequent among highly expressed mouse proteins. **Fig. S4**. Functional analysis of dysregulated proteins defined with a pairwise model in brains of 5xFAD mice. **Fig. S5**. Enrichment analysis of cell-type-specific marker proteins among dysregulated proteins in brains of 5xFAD mice. **Fig. S6**. IPA and gene ontology enrichment analysis of differentially expressed proteins (DEPs) defined with the full model in cortical and hippocampal tissues of 5xFAD mice. **Fig. S7**. Ingenuity pathway analysis (IPA) of Aβ-correlated and anticorrelated DEPs defined by the pairwise model in brains of 5xFAD mice. **Fig. S8**. Strategy to define mouse protein signatures that are concordantly altered also in AD patient brains. **Fig. S9**. Immunofluorescence analysis of 5xFAD brain slices. **Fig. S10**. Analysis of Arl8b protein aggregates using human brain homogenates derived from AD patients and control individuals.

### Additional File 3

**Supplementary Excel File 1.** DEPs from 5xFAD versus wild-type tissue comparisons (hippocampus and cortex); DEPs were defined using a “pairwise model”. **Supplementary Excel File 2.** DEPs in both hippocampal and cortical tissues defined through the “full model”. **Supplementary Excel File 3.** DEPs that correlate or anticorrelate to Aβ aggregate load in both hippocampal and cortical tissues defined through the “pairwise model”. **Supplementary Excel File 4.** Identified genes differentially down- or upregulated (DEGs) in cortex or hippocampus of 5xFAD mice. **Supplementary Excel File 5.** Results of the Ingenuity Pathway Analyses performed in figures 3g and 4e. **Supplementary Excel File 6.** DEPs defined using a “pairwise model” concomitantly altered both in mouse and human brain tissues.

## Declarations

### Ethics approval and consent to participate

All human experiments were performed in accordance with the declaration of Helsinki and approved by Charité University Medicine ethics committee (EA2/118/15), University College London (UCL)/UCL Hospitals Joint Research Ethics Committee and the Ethics Committee of the Capital Region of Denmark (H2-2011-085). Human brain tissue samples for this study were provided by the Newcastle Brain Tissue Resource (NBTR), Newcastle University, UK. All subjects gave written informed consent.

### Consent for publication

Not applicable

### Availability of data and materials

The RNA-seq data have been deposited in the Gene Expression Omnibus (GEO, [99]) database with the identifier GSE198226 (https://www.ncbi.nlm.nih.gov/geo/query/acc.cgi?acc = GSE198226).

The mass spectrometry proteomics data have been deposited in the ProteomeXchange Consortium via the PRIDE [100] partner repository with the dataset identifier PXD030348 (https://www.ebi.ac.uk/pride/archive/projects/PXD030348). The proteomics data are available using the username: and the password: 8Hpbn7JG.

### Authors’ contributions

AB, NN, MK, PS, IB, AS, MZ, CS, LSS, ARW and MR performed the experiments including mass spectrometry and microscopy imaging. AB, CH, EB, AI, ARW, MR, CMM, GD and DB analyzed the data and designed specific experiments. MK, LMB, EJW, JEN, GD, BK and OP provided reagents or technologies. The manuscript was written by EEW with support from AB, CH and SS. EEW acquired the funding and provided overall supervision. All authors read and approved the final manuscript.

## Acknowledgement

We thank Karola Bach, Stephanie Rode, Melanie Humpenöder (MDC Berlin) and Pascal Eede (Charité Berlin) for technical support. We are grateful to the genomics core facility (MDC Berlin) for sequencing RNAs. We thank Lidia Mateo and Patrick Aloy (IRB Barcelona) for providing assistance in data analyses and Thomas Willnow und Anne-Sophie Carlo-Spiewok (MDC Berlin) for providing us breeder pairs of the 5xFAD mouse model. Human brain samples for this study were provided by the Newcastle Brain Tissue Resource which is funded in part by a grant from the UK Medical Research Council (G0400074), by NIHR Newcastle Biomedical Research Centre awarded to the Newcastle upon Tyne NHS Foundation Trust and Newcastle University, and as part of the Brains for Dementia Research Program jointly funded by Alzheimer’s Research UK and Alzheimer’s Society.

## List of abbreviations

FAD: Familial Alzheimer’s disease
Aβ: Amyloid-β
AD: Alzheimer’s disease
MS: Mass spectrometry
LFQ: Label-free quantification
vs.: Versus
CSF: Cerebrospinal fluid
NFTs: Neurofibrillary tangles
wt: Wild-type
tg: Transgenic
IB: Immunoblotting
IS: Immunostaining
MFA: Membrane filter assay
HD: Huntington’s disease
PCR: Polymerase chain reaction
FDR: False discovery rate
DEP: Differentially expressed protein
DEG: Differentially expressed gene
OXPHOS: Oxidative phosphorylation
IPA: Ingenuity pathway analysis
GO: Gene ontology
Q: Quadrant
r: Pearson correlation coefficient
SV: Synaptic vesicle
NfL: Neurofilament light chain
ctrl: Control
SD: Standard deviation
M: Month
H, h: Hippocampus
C, c: Cortex
AU: Arbitrary unit
corr: Correlated
acorr: Anticorrelated
FC: Fold change
Agg: Aggregate

