## Additional File 1 - Suppl Tables S1-S3 for "A proteomics analysis of 5xFAD mouse brain regions reveals the lysosome-associated protein Arl8b as a candidate biomarker for Alzheimer’s disease"

| Name | Supplier | Cat. No. / RRID | Use in this study |
|---|---|---|---|
| Anti-Arl8b | Proteintech | 13049-1-AP / AB_2059000 | IB, IS |
| Anti-Arl8a/b | LSBio | LS-C379507 | IB |
| Anti-β-Amyloid, 1-16 (clone 6E10) | BioLegend | 803002 / AB_2564654 | IB |
| Anti-Lamp1 | Santa Cruz Biotechnology | sc-19992 / AB_2134495 | IB, IS |
| Anti-Calnexin | Stressgen | SPA-860 | IB |
| Anti-VDAC | Cell Signaling Technology | 4661 / AB_10557420 | IB |
| Anti-Golgin97 | Cell Signaling Technology | 13192 / AB_2798144 | IB |
| Anti-Flotillin | Cell Signaling Technology | 3253 / AB_2106734 | IB |
| Anti-NDUFB3 | Abcam | Ab202585 / AB_2890186 | IB |
| Anti-α-Tubulin | Sigma Aldrich | T6074 / AB_477582 | IB |
| Anti-α-Tubulin | Sigma-Aldrich | SAB3501072 | IB |
| Anti-Presenilin-1, NT, clone 2Q127 | United States Biological | P6300-35C | IB |
| Anti-Amyloid β (N) (clone 82E1) | Immuno-Biological Laboratories | JP10326 / AB_2341281 | IS |
| Anti-rabbit IgG peroxidase | Sigma-Aldrich | A0545 / AB_257896 | IB |
| Anti-mouse IgG peroxidase | Sigma-Aldrich | A0168 / AB_257867 | IB |
| Alexa Fluor 594 anti-β-Amyloid, 1-16 (clone 6E10) | Biolegend | 803002 / AB_2564654 | IS |
| Goat anti-Rat IgG (H+L) Cross-Adsorbed Sec. Ab, Alexa Fluor 594 | ThermoFisher Scientific | A-11007 / AB_10561522 | IS |
| Goat anti-Mouse IgG (H+L) Cross-Adsorbed Sec. Ab, Alexa Fluor 594 | ThermoFisher Scientific | A-11005 / AB_2534073 | IS |
| Goat anti-Rabbit IgG (H+L) Cross-Adsorbed Sec. Ab, Alexa Fluor 647 | ThermoFisher Scientific | A-21244 / AB_2535812 | IS |

IS, immunostaining; IB, immunoblotting

**Table S1: Commercial antibodies used in this study**

**AD patients and controls**

| Characteristics | Controls (n = 4) | AD (n = 10) |
|---|---|---|
| *Demographic data* | | |
| Age in years at LP (range) | 67 (60-74) | 76.2 (71-85) |
| Female sex, n (%) | 1 (25%) | 3 (30%) |
| *Clinical data* | | |
| MMSE score at LP?, mean (range), n | 28, n=1 | 22.4 (20-25), n=10 |
| *Biomarker data* | | |
| CSF t-tau pg/ml (range) | 299.25 (239-368) | 814.6 (514-1206) |
| CSF p-tau pg/ml (range) | 39.675 (34-46) | 134.82 (80-225) |
| CSF Aß(1-42) pg/ml (range) | 1277.5 (1021-1508) | 706.1 (388-969) |
| CSF Aß(1-40) pg/ml (range) | 10423.25 (9551-11234) | 11597.1 (6553-15289) |
| CSF Aß(1-42)/Aß(1-40) (range) | 0.122 (0.101-0.139) | 0.062 (0.045-0.077) |

**AD patients and controls**

| Characteristics | Controls (n = 44) | AD (n =38) |
|---|---|---|
| *Demographic data* | | |
| Age in years at LP (range) | 66.16 (47-80) | 75,05 (60-87) |
| Female sex, n (%) | 23 (52.27%) | 20 (52.6%) |
| *Clinical data* | | |
| MMSE score at LP?, mean (range), n | 29.03 (26-30), n=32 | 23.1 (12-30), n=38 |
| *Biomarker data* | | |
| CSF t-tau pg/ml (range) | 268.59 (163-399) | 732.53 (405-1505) |
| CSF p-tau pg/ml (range) | 38.5 (18-57.2) | 111.04 (72-279.2) |
| CSF Aß(1-42) pg/ml (range) | 1170 (724-2110) | 484,5 (229-649) |
| CSF Aß(1-40) pg/ml (range) | 11693.64 (7491-18279) | 10996.82 (4615-18924) |
| CSF Aß(1-42)/Aß(1-40) (range) | 0.0998 (0.059-0.127) | 0,0448 (0.025-0.054) |

**HD patients and controls**

| Characteristics | Controls (n = 10) | HD (n = 10) |
|---|---|---|
| *Demographic data* | | |
| Age in years at LP (range) | 36.6 (27-60) | 46 (25 – 59) |
| Female sex, n (%) | 7 (70%) | 5 (50%) |

**Table S2: Characteristics of AD and HD patients and corresponding controls**

| Biomarker | Spearman correlation coefficient $r_S$ | p value |
|---|---|---|
| Aβ(1-42) | -0.316 | 0.0038 |
| Aβ(1-40) | 0.129 | 0.2466 |
| Aβ42/Aβ40 | -0.405 | 0.0002 |
| t-tau | 0.366 | 0.0007 |
| p-tau | 0.415 | 0.0001 |

**Table S3: Spearman correlation between CSF Arl8b concentration and biomarker values.** The statistical significance of the association was measured with a two-tailed t-test. T-tau, total tau. P-tau, p181 phosphorylated tau.
