## Additional File 2 - Suppl Figures S1-S10 for "A proteomics analysis of 5xFAD mouse brain regions reveals the lysosome-associated protein Arl8b as a candidate biomarker for Alzheimer’s disease"

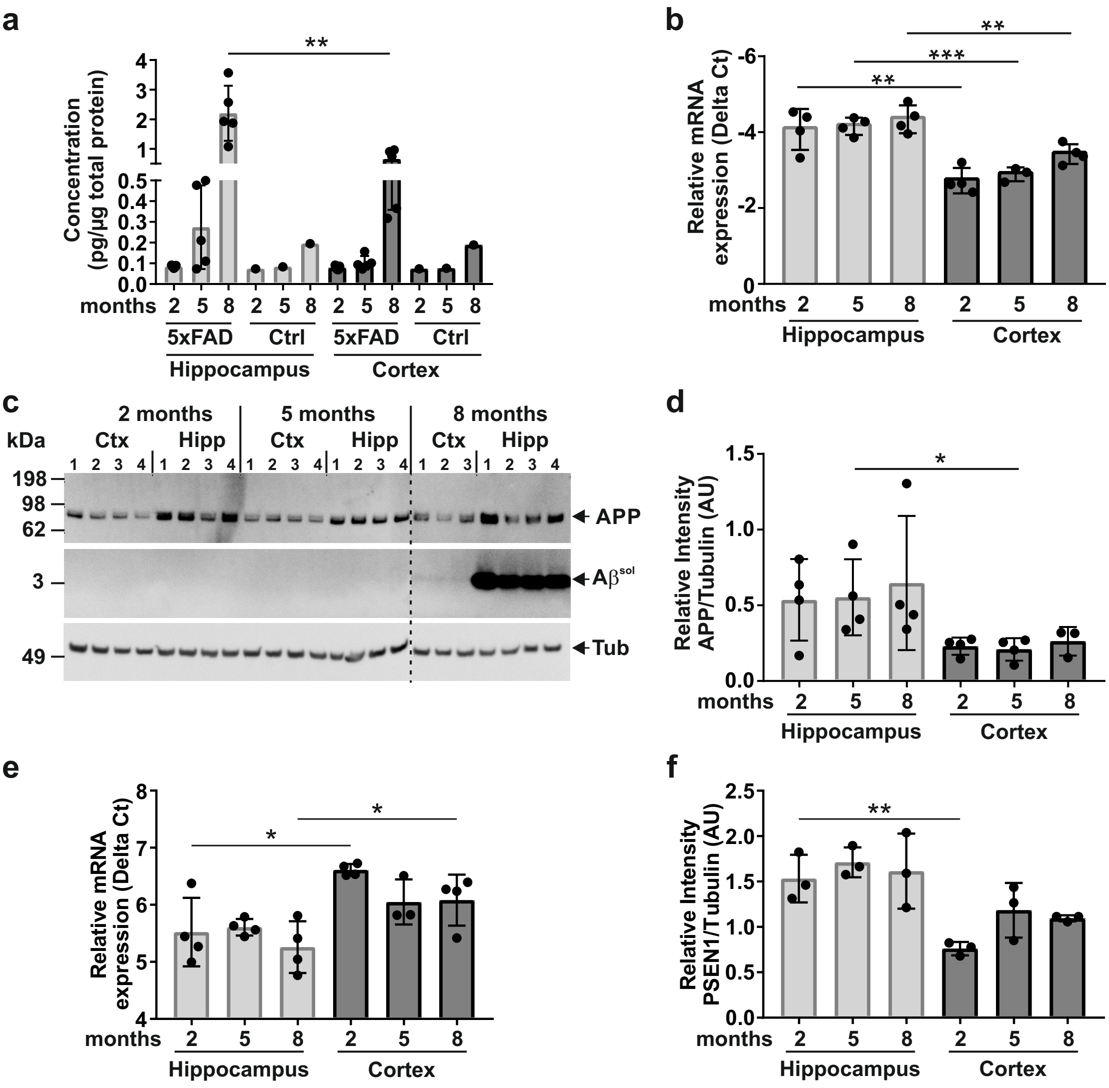


**Figure S1. Aβ peptide levels and APP and PSEN1 expression in hippocampus and cortex of 5xFAD mice.**

**(a)** Hippocampal and cortical brain extracts of aged AD mice (n=5) were used for Aβ40 peptide ELISA. As a control, a pool of wt mouse brain extracts per age and tissue was tested as described in **Figure 1a**. Data represent the mean ± SD of five biological replicates of tg mice per age and tissue. The statistical significance was assessed between hippocampal and cortical tissues of the same age using an unpaired, two-tailed t test (**, p = 0.0077). **(b)** Real Time PCR for quantification of human APP transcript levels in hippocampus and cortex of 2-, 5- and 8-month-old 5xFAD mice. Per age and tissue reverse transcribed cDNAs of four different mice were analyzed using a human APP specific TaqMan gene expression assay. Data were normalized to endogenous mouse EIF-4H and represent the mean ± SD. Statistical analysis was performed using an unpaired two-tailed t test comparing transcript levels of hippocampal and cortical tissues of the same age (2 months Hipp/Ctx: p = 0.0054, 5 months Hipp/Ctx: p = 0.0005, 8 months Hipp/Ctx: p = 0.0065). **(c)** Hippocampal (Hipp) and cortical (Ctx) brain extracts derived from four different 5xFAD mice were analyzed by immunoblotting using 6E10 and anti–α-tubulin (Tub) antibodies (#T6074). Antibody 6E10 recognizes human APP and soluble Aβ ($A\beta^{sol}$). The dashed line indicates that the samples of 8 months old mice run on a separate gel under the same experimental conditions. **(d)** APP and tubulin immunoblot intensities **(c)** were quantified using the Image J. Relative intensity values (mean ± SD) are shown for hippocampal and cortical tissues of 2-, 5- and 8-month-old 5xFAD mice (n=4). Statistical significance was assessed with an unpaired, two-tailed t test (*, p = 0.039). **(e)** Real time PCR for quantification of human PSEN1 transcript levels in hippocampus and cortex of 2-, 5- and 8-month-old 5xFAD mice. Per age and tissue reverse transcribed cDNAs of three to four different mice were analyzed using a human PSEN1 specific TaqMan gene expression assay. Data were normalized to endogenous mouse GAPDH and represent the mean ± SD. Statistical analysis was performed using an unpaired two-tailed t test comparing transcript levels of hippocampal and cortical tissues of the same age (2 months Hipp/Ctx: p = 0.0115, 8 months Hipp/Ctx: p = 0.041). **(f)** Presenilin-1 and alpha-Tubulin (#SAB3501072) expression were quantified by immunobloting using the iBright Analysis Software (Thermo Fisher Scientific). Relative intensity values (mean ± SD) are shown for hippocampal and cortical tissue of 2-, 5- and 8-month-old 5xFAD mice (n=3). Statistical significance was assessed with an unpaired, two-tailed t test (**, p = 0.081).

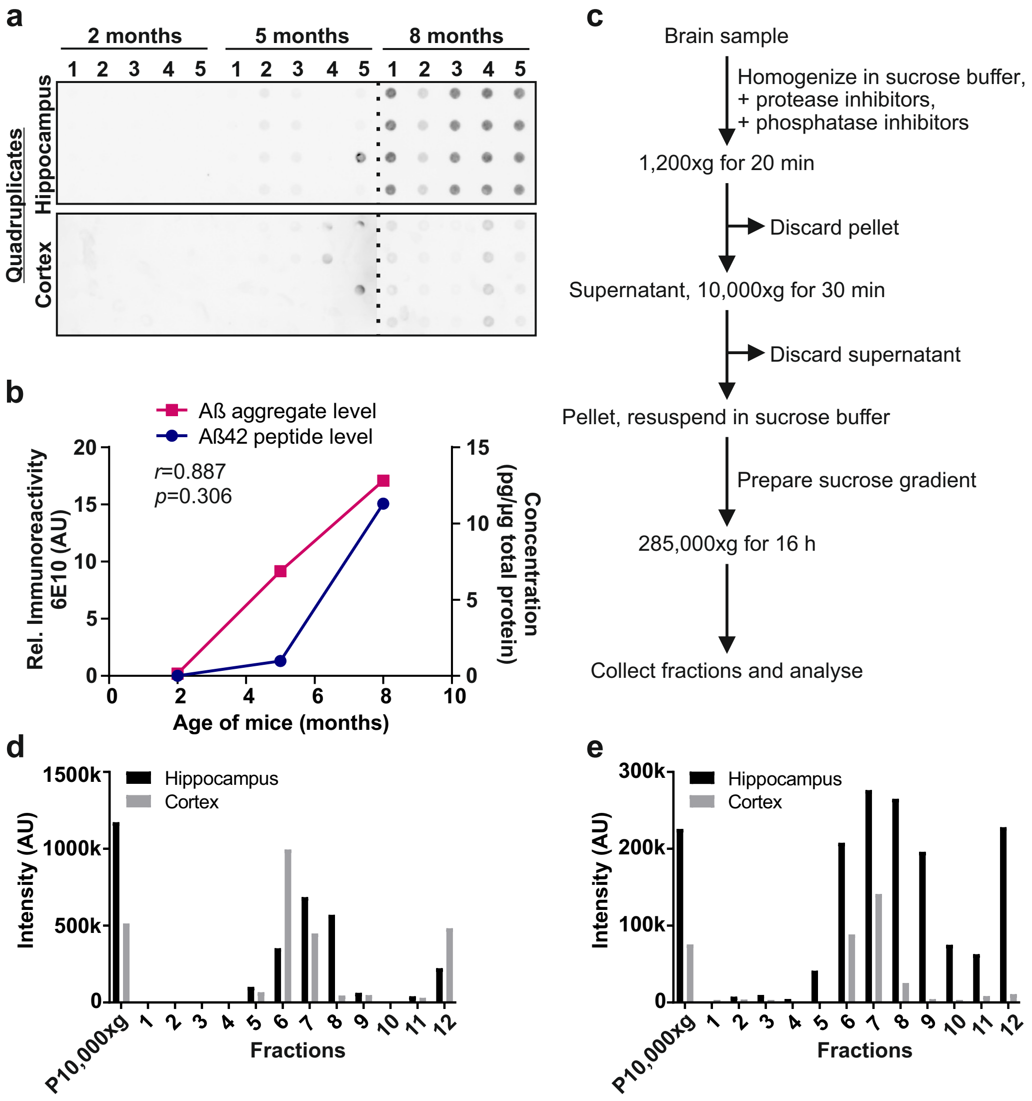


**Figure S2. Analysis of Aβ aggregate formation using membrane filter assays (MFA) and sucrose gradient centrifugations.**
**(a)** Membrane filter assay (MFA) of hippocampal and cortical brain extracts of 2-, 5- and 8-month-old 5xFAD mice. Per age, brain extracts of 5 different mice in quadruplicates were tested. Hippocampal and cortical extracts were analyzed on separate filter membranes. For immunoblotting the antibody 6E10 was used. Both filter membranes were exposed for 1 min. The dashed line indicates where the filter was cut. **(b)** Pearson correlation analysis of Aβ42 peptide levels determined by ELISA (blue line, right axis) and Aβ aggregate levels determined by membrane filter assay (purple line, left axis) in cortical tissue samples. The statistical significance of the association between the Aβ42 peptide levels and Aβ aggregate levels was measured with a two-tailed t-test (not significant, p = 0.306). The correlation coefficient *r* and the p-value are given in the upper left corner of the diagram. **(c)** Schematic representation of membrane fraction purification from mouse hippocampal and cortical brain samples using sucrose gradient centrifugation. **(d, e)** Quantification of soluble APP **(d)** and insoluble Aβ aggregates **(e)** in gel pockets of immunoblots in figure 1g was performed using the iBright Analysis Software (Thermo Fisher Scientific). Immunoblots from hippocampal and cortical fractions for one and the same antibody were prepared under the same conditions.

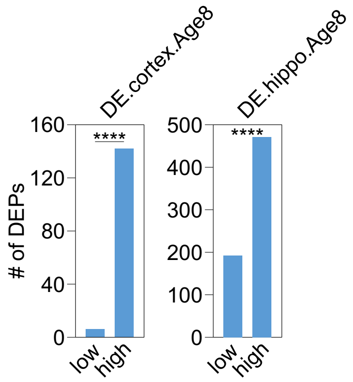


**Figure S3. Analysis of wild-type expression profiles to assess whether protein abundance changes detected in 5xFAD brains are more frequent among highly expressed mouse proteins.** For the differentially altered proteins from DE.Cortex.Age8 and DE.hippo.Age8, a median split was performed and groups of low, medium and high wild-type protein expression were formed, of which the medium group was omitted. The statistical significance was measured with a right-tailed Fisher's exact test (****, $p < 0.0001$). The analysis is based on mean values of measured intensities from five biological replicates of tg mice per age and tissue (n=5).

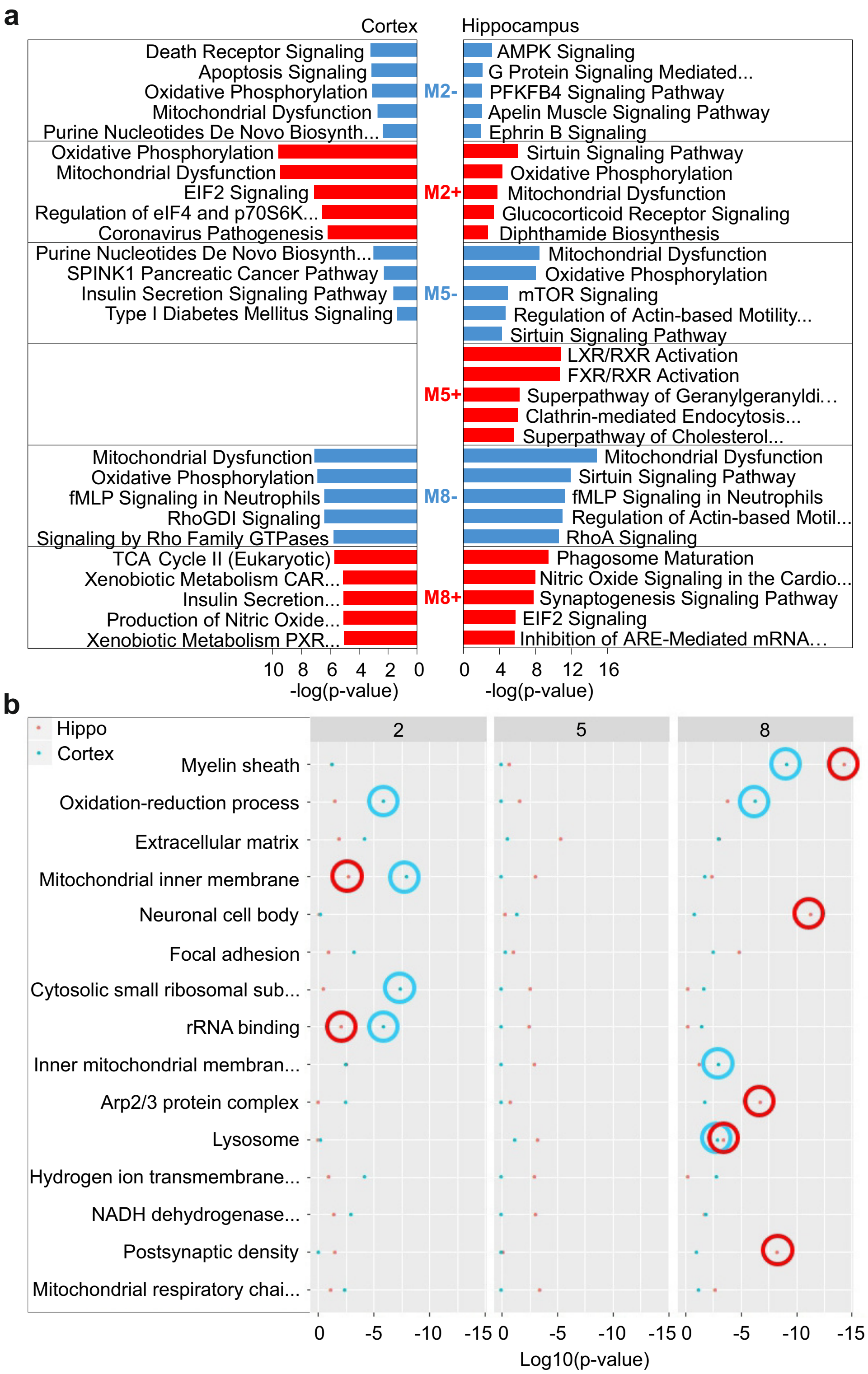


**Figure S4. Functional analysis of dysregulated proteins defined with a pairwise model using data sets from 5xFAD mice and controls.**

**(a)** Ingenuity pathway analysis (IPA) to assess altered cellular processes of dysregulated proteins in hippocampal and cortical tissues; blue indicates downregulated pathways ("-"), red designates upregulated pathways ("+") at 2, 5 and 8 months (M2, M5, M8). The statistical significance was measured with a right-tailed Fisher's exact test to calculate the p-values, adjusted by the Benjamini-Hochberg multiple testing correction. **(b)** Temporal gene ontology (GO) enrichment analysis of DEPs of 5xFAD mouse brains from both hippocampus and cortex at months 2, 5 and 8. Statistical significance was determined using a right-tailed Fisher's exact test to calculate p-values; they are shown as log10 p-values. Highly significant pathways of interest in hippocampus are marked with red circles, in cortex with blue circles. All analyzes are based on mean values of measured intensities from five biological replicates of tg mice per age and tissue (n=5).

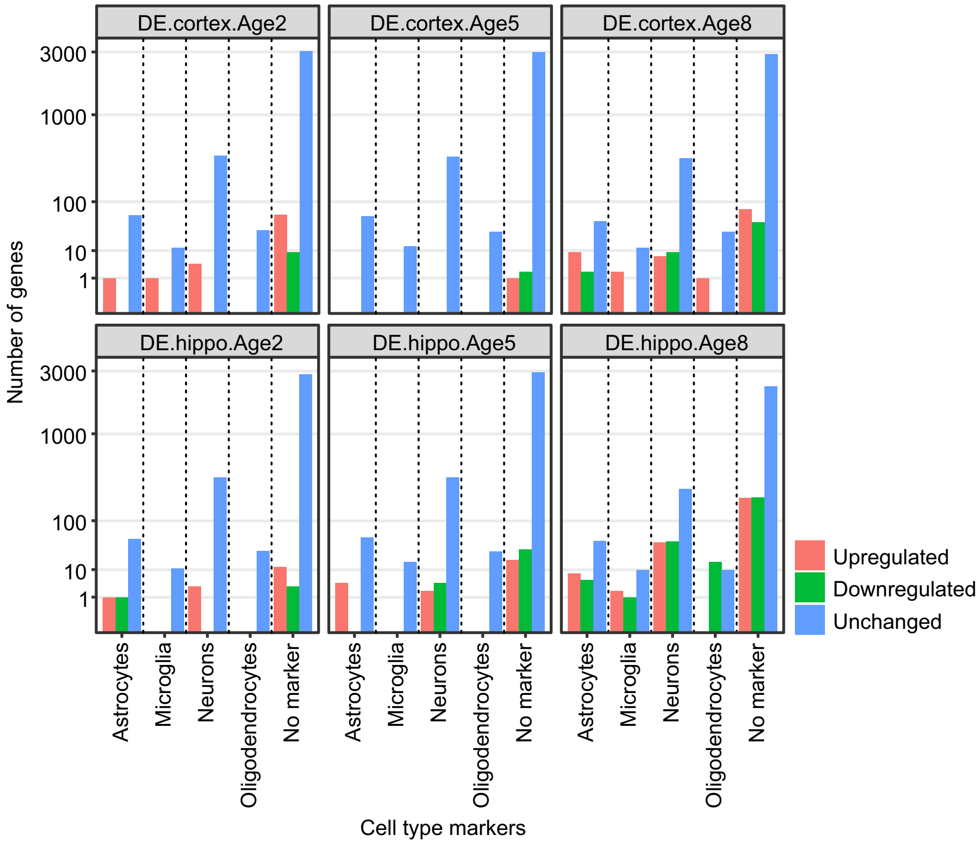


**Figure S5. Enrichment analysis of cell-type-specific marker proteins among dysregulated proteins in brains of 5xFAD mice.**
The numbers of differentially expressed proteins (defined with a pairwise model; y-axis) that are unchanged (blue), down- (green) or up-regulated (red) and enriched in specific cell types (x-axis) are shown. Analyzes are based on mean values of measured intensities from five biological replicates of tg mice per age and tissue (n=5).

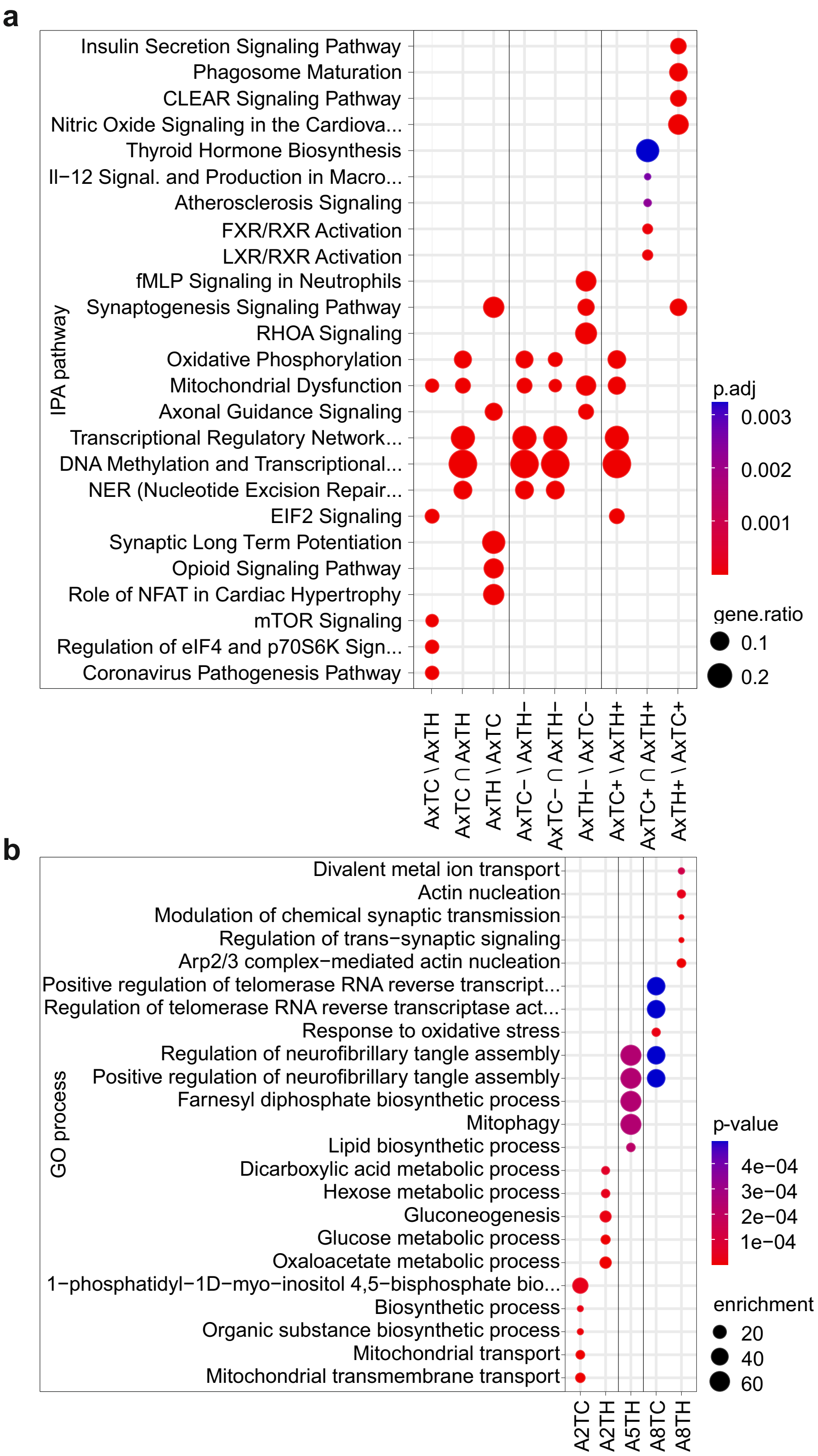


**Figure S6. IPA and gene ontology enrichment analyses of differentially expressed proteins (DEPs) defined with a full model in cortical and hippocampal tissues of 5xFAD mice**
**(a)** Ingenuity pathway analysis (IPA) to assess altered cellular processes of dysregulated proteins in hippocampal and cortical tissues; Groups of dysregulated (left), downregulated ("-", middle) and upregulated proteins ("+", right) across all ages (2, 5 and 8 months) were analysed considering differences ("\") and intersections ("∩") of proteins. The identifiers were denoted analogously as in figures 2a and 2d-f. The statistical significance of the association between the DEPs and the canonical pathway proteins was measured with a right-tailed Fisher's exact test to calculate the p-values, adjusted by the Benjamini-Hochberg multiple testing correction. **(b)** Gene ontology (GO) enrichment analysis of the DEPs identified in hippocampal and cortical tissues of 2-, 5- and 8-month-old 5xFAD tg mice. The enrichment was defined as described in the methods section. Statistical significance was determined by computing an exact p-value of a given minimum hypergeometric (mHG) score, corrected for multiple testing. All analyzes are based on mean values of measured intensities from five biological replicates of tg mice per age and tissue (n=5).

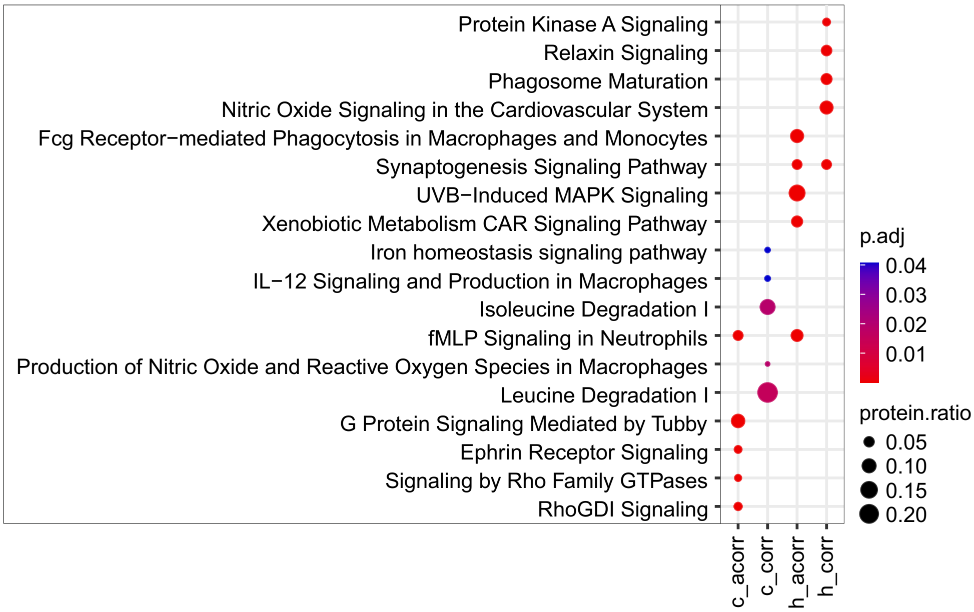


**Figure S7. Ingenuity pathway analysis (IPA) analysis of Aβ-correlated and anticorrelated DEPs defined by the pairwise model in brains of 5xFAD mice.**

Ingenuity pathway analysis with Aβ-correlated and anticorrelated DEPs identified in hippocampal and cortical tissues. The statistical significance of the association between the DEPs and the canonical pathway proteins was measured with a right-tailed Fisher's exact test to calculate the p-values, adjusted by the Benjamini-Hochberg multiple testing correction. All analyses are based on mean values of measured intensities from five biological replicates of tg mice per age and tissue (n=5).

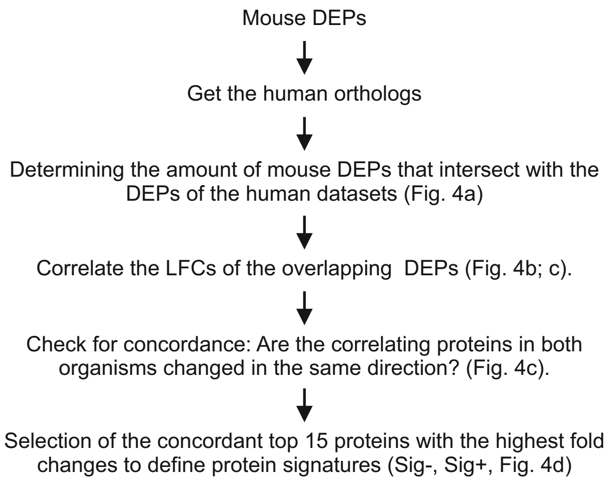


**Figure S8. Strategy to define protein signatures concordantly altered in mouse and AD patient brains.** Schematic representation of the strategy to generate potentially patient-relevant mouse protein signatures (Sig- and Sig+).

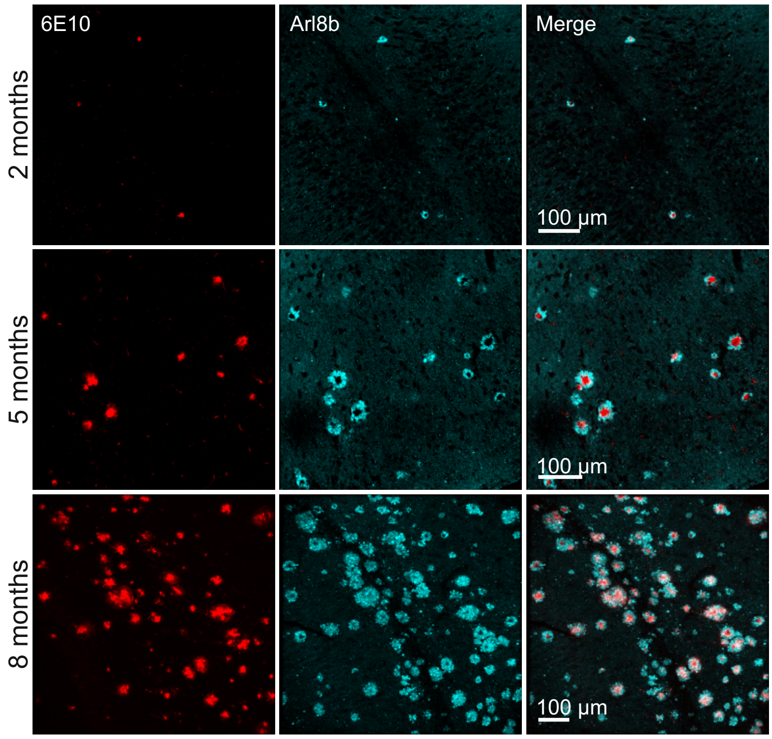


**Figure S9. Immunofluorescence analysis of 5xFAD brain slices.**
Slices of 2-, 5- and 8-month-old mice were stained with AlexaFluor594-labelled 6E10 antibody (red) and anti-Arl8b antibody detected with AlexaFluor647-labelled anti-rabbit IgG (turquoise).

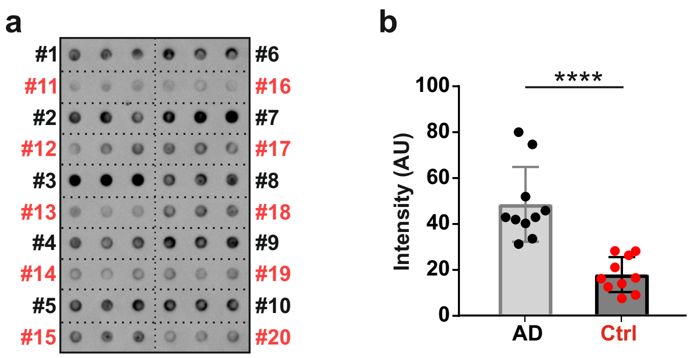


**Figure S10. Analysis of Arl8b protein aggregates using human brain homogenates derived from AD patients and control individuals.**

**(a)** Detection of Arl8b protein aggregates in postmortem brain homogenates of 10 AD patients (1 to 10, black lettering) and 10 age-matched controls (11 to 20, red lettering) using a native MFA. Triplicates per sample were filtered. For immunodetection of Arl8b protein an anti-Arl8b antibody was used. **(b)** Quantification of protein retained on filter membranes was performed using an Aida image analysis software. The statistical significance was assessed with an unpaired, two-tailed t test (****, $p < 0.0001$).
